# A machine learning liver-on-a-chip system for safer drug formulation

**DOI:** 10.1101/2022.09.05.506668

**Authors:** Yunhua Shi, Chih-Hsin Lin, Daniel Reker, Christoph Steiger, Kaitlyn Hess, Joy E. Collins, Siddartha Tamang, Keiko Ishida, Aaron Lopes, Jacob Wainer, Alison M. Hayward, Chad Walesky, Wolfram Goessling, Giovanni Traverso

## Abstract

Drug metabolism leads to biotransformations of pharmaceutical substances that alter drug efficacy, toxicity, as well as drug interactions. Modeling these processes *ex vivo* stands to greatly accelerate our capacity to develop safe and efficacious drugs and formulations. Recognizing the liver as the primary site of drug metabolism, here we report a novel whole-tissue *ex vivo* liver screening platform that enabled modeling of hepatic metabolism and tracking of hepatotoxic drug metabolites. We applied the system for the characterization of acetaminophen (APAP) metabolism and identified interactions that can mitigate the generation of toxic metabolites *ex vivo*. Combining our experimental platform with state-of-the-art machine learning, we validated two novel functional excipients that can prevent APAP hepatotoxicity *in vivo* in mice. To assess translational potential, we prototyped a novel solid dosage form with controlled release of both APAP and our functional excipients. Our this platform provides innovative potential access to actionable data on drug metabolism to support the development of new therapeutic approaches.

## Main

Drug metabolism is a major determinant of the pharmacokinetic and pharmacodynamic properties of administered therapeutic materials^1,2^: through a complex orchestration of metabolic enzymes, the active pharmaceutical ingredient (API) will undergo a series of chemical transformations that can substantially alter the physicochemical and pharmacological properties of a drug. This can, for example, lead to the activation or inactivation of a drug^3,4^ or create toxic species^5,6^. A systematic understanding of a drug’s metabolic profile is crucial to ensuring that therapeutics are clinically efficacious and safe. Accordingly, a multitude of experimental and computational approaches have been developed that anticipate the enzymes involved in the metabolism of a pharmaceutical compound^7^. Although the enzyme-substrate relationships and the resulting chemical transformations can often be predicted *in silico* and *in vitro*, the complex and dynamic environment found in physiological conditions can behave very differently from simplified model systems and cause unexpected clinical effects^2^: physiologically relevant cell composition, ratio and arrangement, enzymatic expression levels and physical barriers are often inaccurately captured in simplified *in silico* or *in vitro* models^8^. Increasingly complex “organ-on-a-chip” systems are engineered to model physiologically relevant factors such as complex compositions of various cell types, physical forces, or pH and oxygen gradients, and have enabled advances in reproducing some physiological effects by approximating the *in vivo* conditions for specific applications^9^. However, such engineered systems are inherently driven by the current mechanistic understanding of the respective microenvironment and therefore might inaccurately capture less well understood or more complex physiological processes *in vivo*. The development of platforms that accurately represent physiological conditions while enabling high-throughput screening and perturbations remains a research area of high translational significance.

Even in cases where the metabolic pathways of a drug are well understood, the metabolism of multiple drugs across such pathways can cause competitive inhibitions that dramatically shift compound metabolism and lead to unexpected and often adverse clinical outcomes^6,10^. These effects are the root cause of drug-drug interactions, which can inactivate life-saving therapeutics or result in drug overexposure with potentially fatal outcomes^11^. Significant scientific and clinical resources are devoted to identifying potential drug-drug interactions in order to avoid adverse effects on patients. More recently, drug interactions have been harnessed to alter metabolism in a controlled manner in order to improve the pharmacokinetic properties of a difficult-to-deliver therapeutic: functional formulations can enable the delivery of life-saving therapeutics that would otherwise have unacceptable metabolic profiles. In this regard, it is becoming apparent that chemicals considered safe for human consumption at common dosages, such as excipients and food additives, can affect drug pharmacokinetic activity that can therefore cause excipient-drug or food-drug interactions^12-16^. Such materials provide ideal starting points for the design of novel functional formulations since they can often be readily deployed in novel formulations with accelerated approval mechanisms.

Here, we report the development of a platform that combines biological data from a novel whole-tissue system with state-of-the-art machine learning to enable the rapid identification of functional formulations with high translational potential. As a proof of concept, we created and validated a new *ex vivo* liver model that enabled us to predict the hepatic metabolism and associated hepatotoxicity of acetaminophen. APAP is the most widely used over-the-counter analgesic and antipyretic in Europe (marketed as paracetamol) and the USA^17^ and occurs twice on the World Health Organization’s (WHO) list of essential medicines for the treatment of pain or acute migraine attacks^18^. However, APAP is also recognized to be associated with severe hepatotoxicity, for example when exceeding recommended dosages, when taken by patients with pre-existing liver conditions, or in combination with alcohol intake or fasting^19^. These challenges are further exacerbated through increasing usage of APAP as well as availability of high-dose formulations^20^. In the US alone, APAP overdose leads to 30,000 hospitalizations annually^21^ and it has been specifically associated with adverse outcomes in vulnerable populations. The associated liver toxicity originates from the cytochrome P450 metabolite N-acetyl-p-benzoquinone imine (NAPQI), a toxic byproduct which causes an elevation in mitochondrial reactive oxygen species (ROS), DNA damage, and liver necrosis ^22^. Despite this mechanistic understanding, APAP toxicity is the most common cause of liver transplantation in the US and causes at least 500 US deaths per year^23^. Novel APAP formulations that modulate its metabolism could mitigate such adverse effects to provide safer and more efficacious medications. Using publicly available biochemical data, we generated *in silico* predictive models of cytochrome P450 inhibition in order to identify novel modulators of APAP metabolism. Our pipeline successfully identified two excipients as previously unknown inhibitors of cytochrome P450 enzymes, and we validated their capacity to prevent acetaminophen-associated hepatotoxicity *in vivo*. Finally, we prototyped a novel APAP dosage form with a carefully designed release profile of our protective agent. Taken together, this proof of concept demonstrates the capacity of our platform and extensions thereof to facilitate the design of safer and more effective therapeutic candidates.

## Results

### *Ex vivo* porcine liver tissue accurately captures hepatotoxic effects of APAP metabolites

Porcine liver tissue is an ideal model for studying drug metabolism given its physiological similarity to human liver tissue^24^. We investigated the homology of the ten major cytochrome P450 enzymes between humans and pigs and found a >75% and a >80% conservation in amino acid and mRNA respectively (NCBI; *cf*. Figure S1). We have previously demonstrated the ability to *ex vivo* cultivate porcine intestinal tissue while retaining key physiological properties for screening^25^; here, our goal was to develop a high-throughput system that could facilitate interrogation of hepatocyte response in the context of native organ architecture. The complex structure of the liver required additional processing in order to produce tissue samples that could be integrated into customized plates (Figure 1A)^25^. Briefly, we used a dermatome to produce consistent 250 μm thin layers of intact liver tissue from fresh procured porcine liver. Such thin layers enabled us to retain the microenvironment of the intact liver tissue while simplifying and standardizing the application of biological or chemical perturbations and the assessment of various tissue properties. For example, by retaining a similar number of cells across different tissue patches (Figure S2), we are able to calculate relative viability. Specifically, we used a standard, commercially-available assay that measures ATP levels in order to quantify the viability of the procured porcine liver tissue over time. We confirmed that our tissue retained high viability (77 ± 32% compared to fresh tissue) for up to 20h (Figure 1B) and retained key biological characteristics. We hypothesized that this retention would enable us to investigate more complex phenotypic phenomena such as the cytotoxic effects of certain treatments like acetaminophen. To this end, we treated the liver slices with increasing doses of APAP and measured tissue viability with our ATP-based assay. We found a dose-dependent effect on the liver tissue, with severe tissue damage and necrosis observed when the APAP concentration reached 10 mM (Figure 1C). To further assess the resolution of this assay system, we investigated the effect of N-acetyl-meta-aminophenol (AMAP), a metabolite with reduced toxicity potential; as expected, AMAP treatment resulted in higher viability compared to APAP treatment (Figure S3). We validated that our system was able to capture hepatotoxic effects of other drugs such as chlorozoxazone and diclofenac (Figure S4). Taken together, these results indicate that our liver tissue slices maintained the ability to metabolize drug compounds and react to toxic species in a manner that closely resembles physiological conditions.

**Figure 1.**
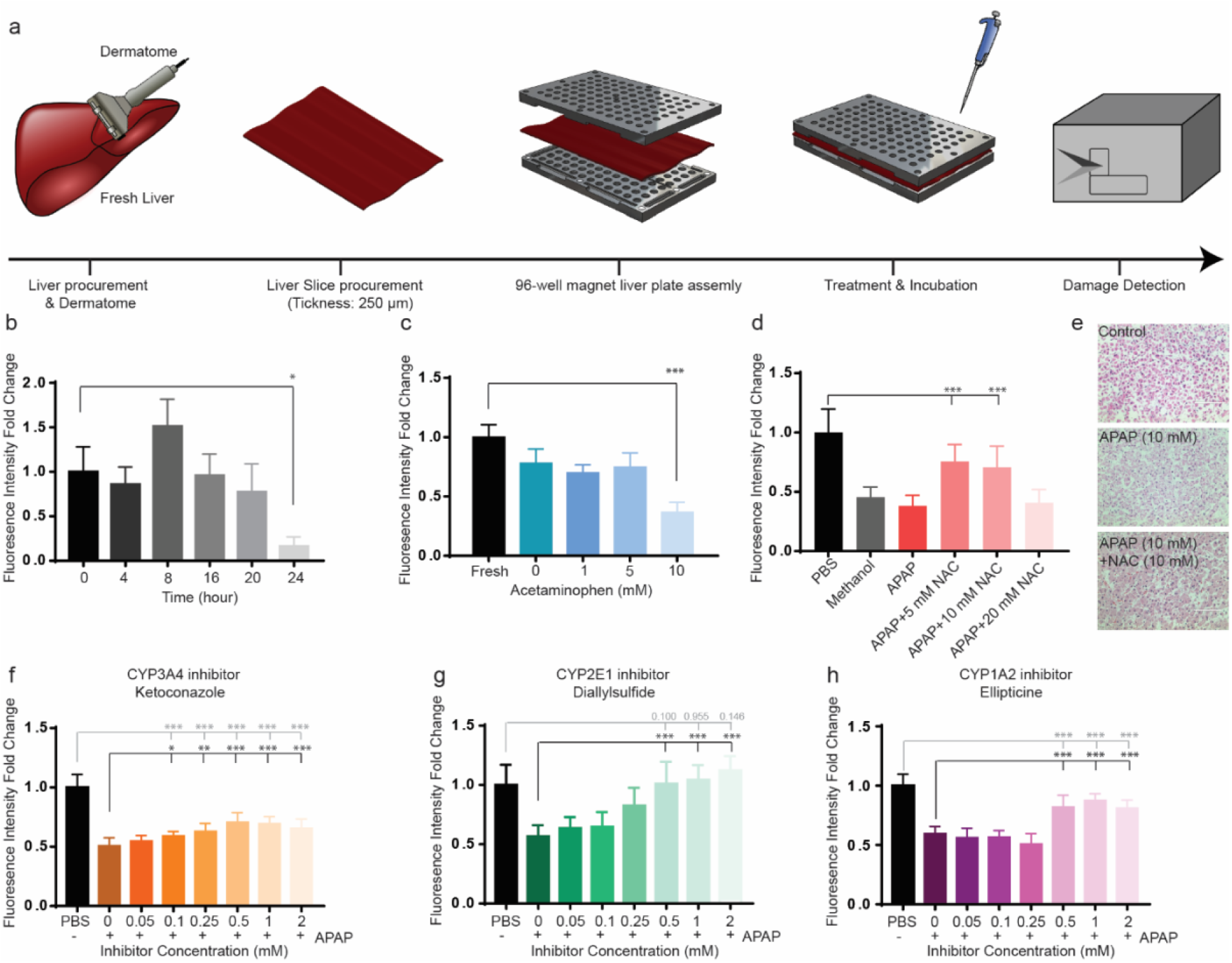
Validation of the *ex vivo* liver-on-a-chip system. **A**) Schematic of the *ex vivo* tissue procurement and platform assembly protocol. **B**) Viability of the *ex vivo* cultured liver tissue. Tissue slices were incubated in culture medium for different durations. Viability was determined using an ATP-based Live/Dead assay as described in Materials and Methods. Acquired viability data was normalized to quantify relative tissue viability compared to freshly procured tissue slices. **C**) Cytoxicity of APAP treatment. The *ex vivo* liver slices were treated with various concentrations of APAP and viability of the treated tissue was calculated using the ATP-based Live/Dead assay. Acquired viability data was normalized to quantify relative tissue viability compared to the PBS-treated control group. **D**) Pretreatment with NAC, a well-known hepatoprotective agent reversing acute liver damage caused by APAP overdose. NAC pre-treatment at various concentrations was tested for its ability to improve relative tissue viability after APAP treatment. **E**) H&E staining of the *ex vivo* tissue previously treated with APAP and/or NAC visualizes reversible tissue damage. **F)-H)** Reversal of hepatotoxicity of APAP through known inhibitors of CYP3A4 (ketoconazole, F), 2E1 (diallyl sulfide, G), and 1A2 (ellipticine, H). Various concentrations of inhibitors were tested to prevent tissue damage when co-administered with 10 mM APAP and relative viability was determined through the ATP-based Live/Dead assay. For the *ex vivo* study, n=3 animals are included for each screening, and a parallel of m>10 is conducted on the same animals for each condition. Therefore, a total of >30 repeats in included. Data analysis is processed as described in “Statistical analysis” of Material and Method.

#### Inhibition of cytochrome metabolism reverses APAP toxicity ex vivo

After validating that our platform could be used to measure the differential hepatotoxic potential of drug metabolites, we investigated whether known drug interactions that reduce adverse metabolism and associated toxic species can be reproduced on our platform. N-acetyl-l-cysteine (NAC) is well known to prevent acute liver damage caused by APAP overdose *in vivo*^26^. Using our system, we co-treated porcine liver tissue with 5 mM or 10 mM of NAC and found that it effectively prevented APAP toxicity and fully rescued tissue viability to control levels (Figure 1D & E). However, higher concentrations of 20 mM NAC did not provide protection (Figure 1D and S3a), which is potentially caused by the toxicity associated with high concentrations of NAC^5,27^. Recognizing that cytochrome metabolism is responsible for transforming APAP into hepatotoxic species, we next investigated whether our platform can capture protective effects of other cytochrome inhibitors. We selected three drugs with known cytochrome inhibitory potential (diallyl sulfide, ketoconazole, and ellipticine) and tested their ability to reverse APAP toxicity *ex vivo*. All three inhibitors protected the tissue from APAP toxicity (Figure 1F-H); in particular, tissue treated with diallyl sulfide was fully protected (Figure 1G). Diallyl sulfide also provided protection from cytotoxic AMAP metabolites (Figure S3d). Ketoconazole and ellipticine provided only partial protection; this could be due lower inhibitory potency against cytochrome P450 enzymes or because the inhibitors themselves exert cytotoxic effects. Accordingly, combinations of these inhibitors at lower concentrations resulted in improved protective effects compared to treatment with a single inhibitor (Figure S3). Similarly, we were able to validate that inhibition of the cytochrome P450s 3a4, 2c9 or 2e1 could mitigate hepatoxicity caused by the drugs diclofenac and chlorzoxazone. This data confirmed that by inhibiting various cytochrome enzymes, the hepatotoxicity caused by different drug metabolites could be reversed in our *ex vivo* liver tissue.

### Machine learning identifies rutin and eugenol as cytochrome inhibitors with protective effects against APAP toxicity *ex vivo*

Our initial experiments with known cytochrome inhibitors were consistent with previous findings, which attests to the ability of our model to capture the protective effects of these agents on liver tissue. However, there is a need to identify novel protective agents. Many of our investigated agents are potent drugs themselves and have adverse effect profiles. Indeed, we had observed that several of these agents caused cytotoxicity at elevated concentrations (Figure 1), which has important consequences for clinical translation. Furthermore, many of the investigated agents did not fully protect against APAP toxicity, which indicates that the inhibitory effects are weak; full protection would require a cocktail of multiple protective agents, increasing the risk of unknown drug interactions (Figure 1). Therefore, we decided to combine our acquired data with publicly available cytochrome P450 inhibition data to accelerate the identification of novel protective agents through machine learning.

By combining data from our experiments, the literature, and the ChEMBL24 database^28^, we generated a dataset with a total of 50,552 data points that captures the molecular signatures necessary to inhibit the three cytochrome enzymes involved in the metabolism of acetaminophen, cytochrome P450 1a2, 2e1, and 3a4. In our previous research, we have established random forest-based machine learning models in order to identify functional excipients that were able to modulate metabolism and transport of specific therapeutics^16^; here, we adapted this technique in order to identify new cytochrome metabolism modulators (Figure 2A). Our model was based on highly heterogeneous data, including a variety of different experimental protocols. As a result, the model’s retrospective accuracy values were low, since cross-validation inappropriately prioritizes different mechanisms and readouts in the training. However, since our goal was to prioritize novel cytochrome inhibitors among the top predicted candidates, we estimated BEDROC enrichment values. This analysis showed that, when sorting predictions according to the associated predictive confidence, the model successfully ranked inhibitors of P450 1a2, 2e1, and 3a4 higher than non-inhibitors among the top 1% most confident predictions.

**Figure 2.**
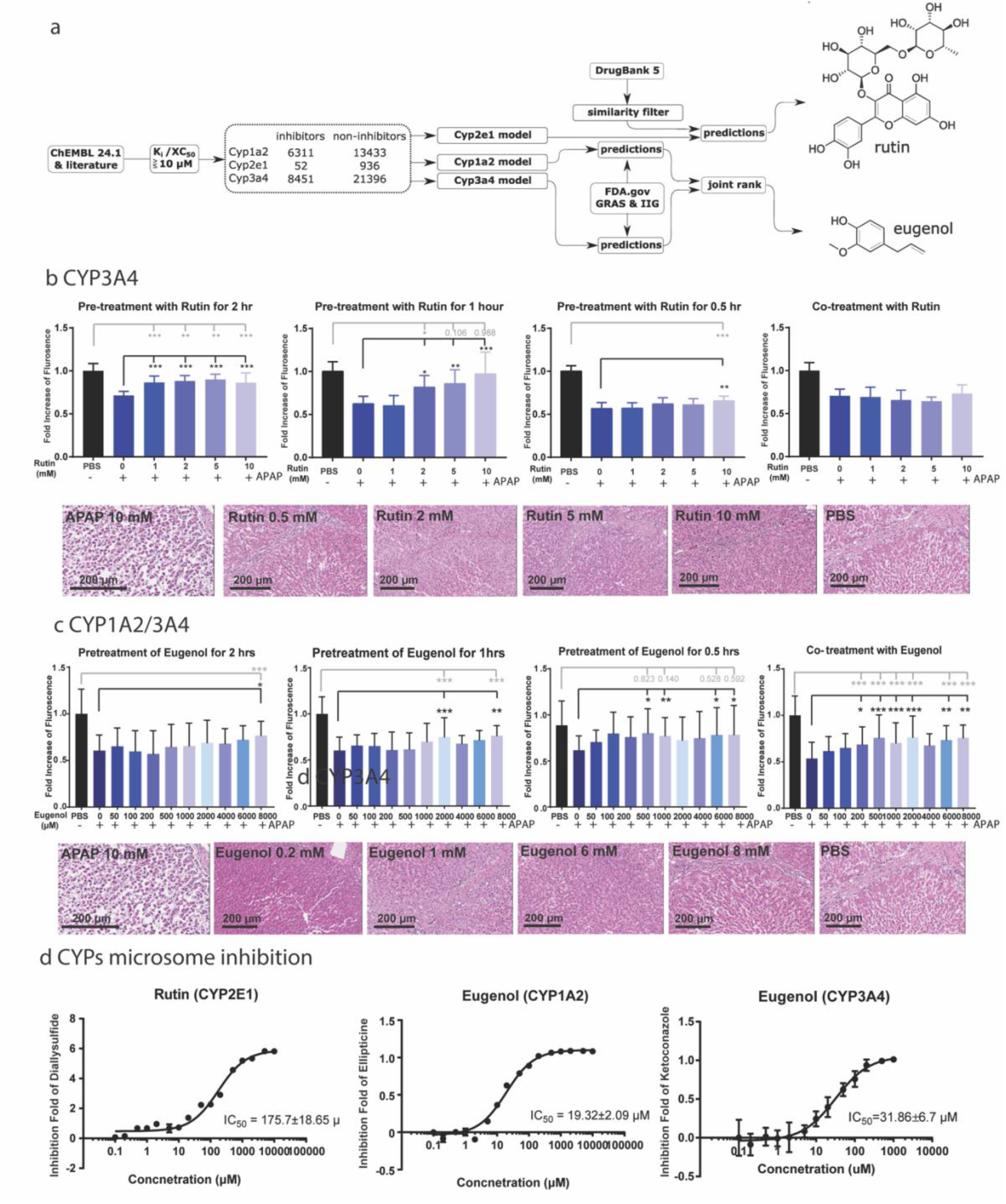
Machine learning identified novel cytochrome inhibitors that prevent APAP-induced tissue damage *ex vivo*. **A**) Schematic of *in silico* workflow for the identification of new cytochrome inhibitors and chemical structures of hits rutin and eugenol. **B**) Rutin reduced APAP liver hepatotoxicity in a concentration and time dependent manner (n=3 animals and m>10, a total of >30 repeats). **C**) Rutin reduced APAP liver hepatotoxicity in a concentration and time dependent manner (n=3 animals and m>10, a total of >30 repeats). **D**) A human microsome inhibition assay was conducted to determine the functional effect of rutin on CYP3A4 and eugenol on CYP1A2 and 3A4. Each concentration is conducted with 12 repeats and the kinetic curve is fitted into a Dose-response-Inhibition function to obtain IC_50_. In part B and C, the data is analyzed with an ANOVA model, and a value of p < 0.05 was considered statistically significant (* for p < 0.05, ** for p < 0.01 and *** for p < 0.001).

We first attempted to identify novel cytochrome P45 2e1 inhibitors from the chemical structure of approved, small molecular compounds included in the DrugBank 5.0 collection. To ensure novelty from previously published cytochrome modulators, we excluded compounds that had similar substructures to the known 2e1 inhibitors in our training database (Tanimoto similarity < 0.5 based on Morgan fingerprints with radius 2 using 2048 bits). We predicted 2e1 inhibition for each of these compounds and then sorted them according to predictive confidence (i.e., maximum number of trees in the forest predicting a modulation). Our model predicted that rutin and suramin, respectively, would be the top two cytochrome 2e1 inhibitors. In our *ex vivo* porcine tissue model, we varied the timing and dose of both rutin and suramin in order to understand their protective potential under different treatment protocols. When administered one hour before APAP exposure, suramin provided only partial protection at doses larger than 100 μM (Supplementary Material). Rutin fully protected the liver against APAP toxicity (Figure 2B). To confirm that the protection was due to cytochrome 2e1 inhibition, we conducted a microsome-based, *in vitro* functional assay; we found that rutin inhibits cytochrome P450 2e1 in a dose-dependent manner with an IC_50_ value of 175.7 ± 18.65 μM.

To identify novel protective agents with higher efficacy, we combined our computational models for 1a2 and 3a4 inhibition. Our goal was to identify dual-inhibitors of these two enzymes among Generally Recognized As Safe (GRAS) and Inactive Ingredient (IIG) substances. To do this, we used our computational model to predict the inhibitory potential of all compounds in these libraries; the predictive confidence was then used to sort candidates according to their average rank for both proteins. We eliminated all chemical dyes from the list given both their allergenic potential^15^ and the generally low doses that these agents are approved for, which would hamper their potential to be used in functional formulations. From this list, the two top predictions were eugenol and cyanocobalamin. In our *ex vivo* porcine liver model, cyanocobalamin did not provide any protection under the investigated conditions; however, eugenol was able to reduce APAP toxicity at various conditions, including when it was co-administered with APAP (Figure 2C). We tested whether eugenol indeed inhibited cytochrome 1a2 and 3a4 in our functional *in vitro* assay and found a dose-dependent inhibition against both enzymes (Figure 2D). Notably, eugenol was part of our original training data but was annotated as inactive through the DrugMatrix screen because it exhibited low (<50%) inhibitory activity at a testing concentration of 10 μM^29^. The random forest model was able to contextualize this data and predict that eugenol could be a promising agent for our purposes at higher supplemented concentrations despite this negative annotation.

To assess whether eugenol and rutin use complementary mechanisms to exert their protective effects, we co-administered the two excipients *ex vivo* (Supplementary Figure 5). We found that the protective effect was indeed enhanced in a non-competitive manner (Supplementary Figure 14). This demonstrates that complex functional formulations with multiple protective agents can be designed to reduce hepatoxicity risks.

### Rutin and eugenol prevent APAP toxicity *in vivo* without disrupting other metabolic pathways

To test whether rutin and eugenol would provide protection from APAP toxicity *in vivo*, we modeled APAP overdose in BALB/c mice. These mice develop severe acute liver damage after a single dose of 350 mg/kg APAP *via* intraperitoneal injection^30,31^. The *ex viv*o experiments indicated that the protection rutin and eugenol provided was time-dependent, with rutin providing the best protection when pre-incubated for 1h while eugenol performed best with 0.5h pre-incubation. However, we recognized that *in vivo* distribution and metabolism might alter exposure dynamics and therefore hinder direct translation of the *ex vivo* results. Therefore, we explored different treatment regimens that dosed rutin and eugenol at different time points before APAP administration. Similarly, we also explored the effect of using a range of different concentrations of rutin and eugenol. As our experimental readout, we quantified serum-levels of alanine transaminase (ALT) and aspartate transaminase (AST); two commonly used markers of hepatotoxicity.

A co-treatment or 30-minute pre-treatment with rutin in a range of 20-80 mg of rutin/kg animal weight significantly protected the liver from APAP toxicity *in vivo* (Figure 3A & 4B). Full protection was achieved at rutin doses below 80 mg/kg. Longer pre-treatment periods (1 or 2 hours) did not provide protection. Eugenol provided protection when a concentration of 40-120 mg per kg of animal weight was co-administered with APAP or when 120 mg/kg was given 30 minutes prior to APAP administration (Figure 3C & 4D). These promising results demonstrate that the two identified functional excipients can prevent hepatoxicity *in vivo;* however, the difference in dosages and timing from our *ex vivo* experiments indicate that the absorption dynamics of rutin and eugenol need to be considered when translating the *ex vivo* findings.

**Figure 3.**
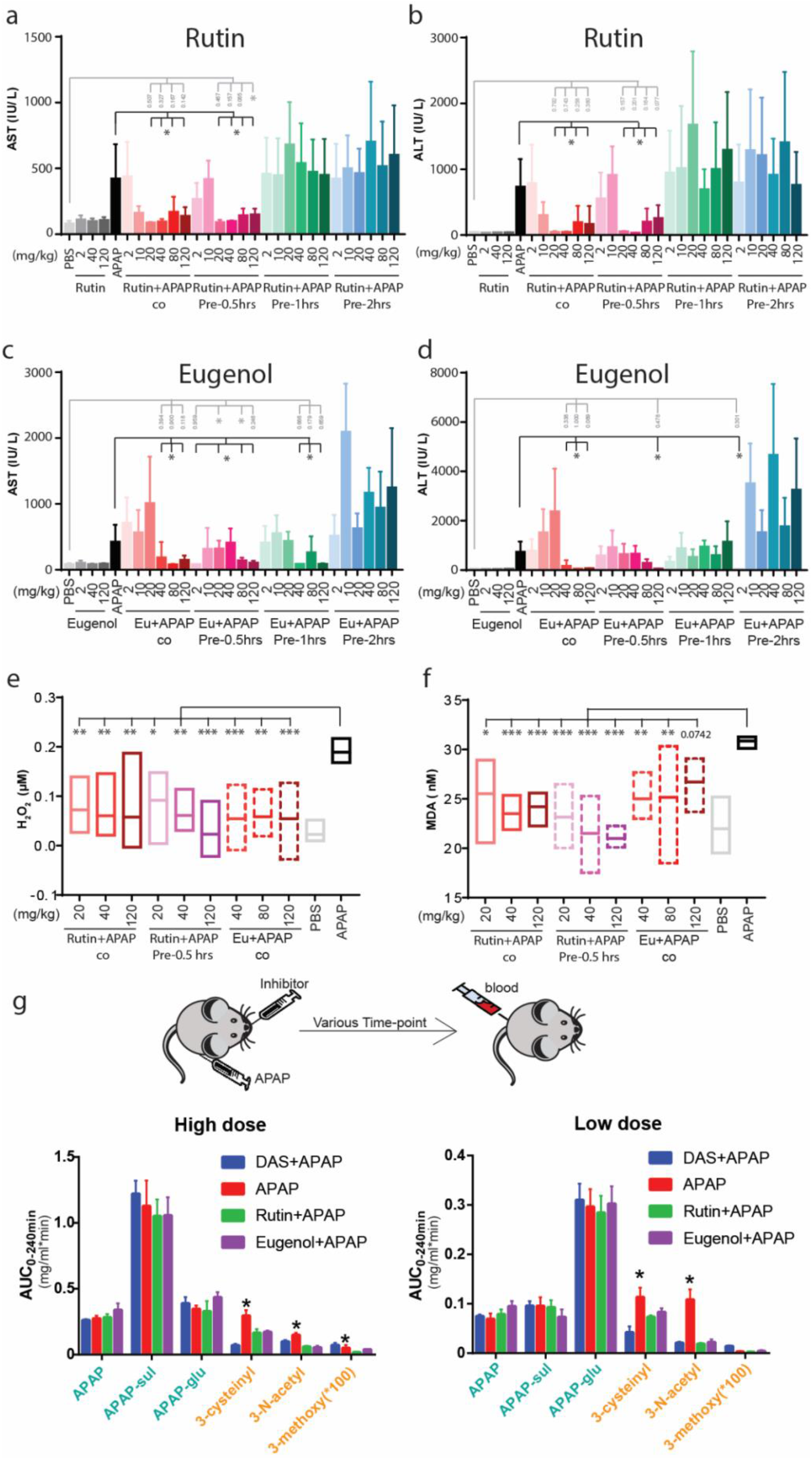
In vivo validation of hepatoprotective effects of rutin and eugenol. **A**) AST levels of mice treated with different concentrations of rutin when administrated 2 h, 1 h, 0.5 h or co-administrated with a high APAP dose. Rutin was delivered through oral gavage, while APAP was delivered via I.P. injection. **B**) ALT level detection of same mice from panel A. **C**) AST levels of mice treated with different concentrations of eugenol when administrated 2 h, 1 h, 0.5 h or co-administrated with APAP. Eugenol was delivered through oral gavage, while APAP was delivered via I.P. injection. **D**) ALT level detection of same mice from panel C. **E**) H_2_O_2_ level of APA-induced liver damage and reduction through rutin or eugenol. **F**) MDA level indicating mitochondria oxidation through APAP overdose and protection from this effect by rutin or eugenol. **G)** In-depth analysis of presence of various APAP metabolites after a high- or low-dose APAP treatment or co-treatment of a high- or low dose of APAP with rutin, eugenol or DAS. Serum concentrations of toxic metabolites (orange) and safe metabolites (green) were recorded at various times to determine AUC values as measures of total concentration. While safe metabolites were not affected, toxic metabolites were mitigated. For the *in vivo* study, a total of 5 mice for each data point is included in this study. Method of data analysis is described in material and method section in detail.

To further validate that our identified functional excipients would provide protection using the developed treatment protocol, we included other readouts of liver damage. APAP overdose has also been associated with mitochondrial oxidation damage^22,32^. To investigate whether rutin and eugenol could prevent mitochondrial oxidation, we measured levels of hydrogen peroxide (H_2_O_2_) and Malondialdehyde (MDA), two biomarkers of oxidative stress, in the mice that received fully protective formulations. APAP overdose alone resulted in an increase in both H_2_O_2_ and MDA; however, in all treatments with rutin or eugenol, there was a significant reduction in both markers (Figure 3E&F).

The specific metabolites responsible for APAP hepatotoxicity are NAPQI and its toxic byproducts, 3-cysteinyl acetaminophen, 3-N-acetyl-cystein-s-yl-acetaminophen, and APAP glutathione^33^. In order to measure whether our newly identified formulations reduced these toxic metabolites, we administered a “normal” dose (100 mg/kg) as well as an “overdose” (350 mg/kg) of APAP to mice and tracked the plasma levels of the APAP metabolites during a 4-hour therapeutic window. For the normal dose and the overdose, both rutin and eugenol significantly reduced the area under the curve (AUC) of all toxic species, while plasma levels of APAP itself and its benign metabolites (APAP sulfate and APAP glucuronide) were not altered (Figure 3 G&H). These *in vivo* results indicate that both rutin and eugenol provide protection from APAP overdose-associated hepatotoxicity by reducing the toxic, oxidizing metabolites that damage mitochondria and liver cells without altering the pharmacokinetics of APAP itself.

Finally, we wanted to determine whether other mechanisms instead of cytochrome inhibition could be responsible for the observed protective effects of our formulations, such as an increase in glutathione (GSH) generation or a reduction of cytochrome P450 enzymes expression. We performed a total GSH analysis and a quantitative polymerase chain reaction (qPCR) of liver tissue extracted from the treatment groups (*cf*. Figure 2). We found that the rutin- and eugenol-treated groups did not have elevated levels of GSH in their liver tissue compared to the control group. Notably, APAP alone depleted GSH, which was most likely caused by the increased production of NAPQI (Figure S11). Furthermore, none of the expression levels of any of the major cytochrome P450 enzymes were affected by the administration of rutin, eugenol, or APAP (Figure S12).

### Solid dosage forms can release APAP and functional excipients

We have shown that our functional excipients are most effective at preventing hepatotoxicity when they are delivered prior to APAP exposure. However, an ideal solid dosage form would enable the oral co-administration of APAP together with these protective agents to avoid complex multi-pill dosing regimens. Our goal was to develop a dosage form that would release rutin or eugenol prior to the release of APAP. We designed a pill with two distinct hemispheres: one contained APAPand the other contained rutin (Figure 4A). We formulated the two parts independently, with each part designed to have different release kinetics: APAP was mixed with carboxyl cellulose/magnesium stearate and rutin was mixed with sodium bicarbonate/citric acid. The APAP mixture was designed to have a slower degradation rate than the rutin mixture. The two individual formulations were pressed layer-by-layer under high pressure (26,000 N) to form a single pill. *In vitro* release studies in simulated gastric fluid (SGF) confirmed that our layered formulation released rutin approximately 15 minutes before the APAP was released (Figure 4B). This proof of concept demonstrates the feasibility of designing safer APAP pills that pre-release our functional excipients for potentially safer clinical administration of important therapeutics with less side-effects.

**Figure 4.**
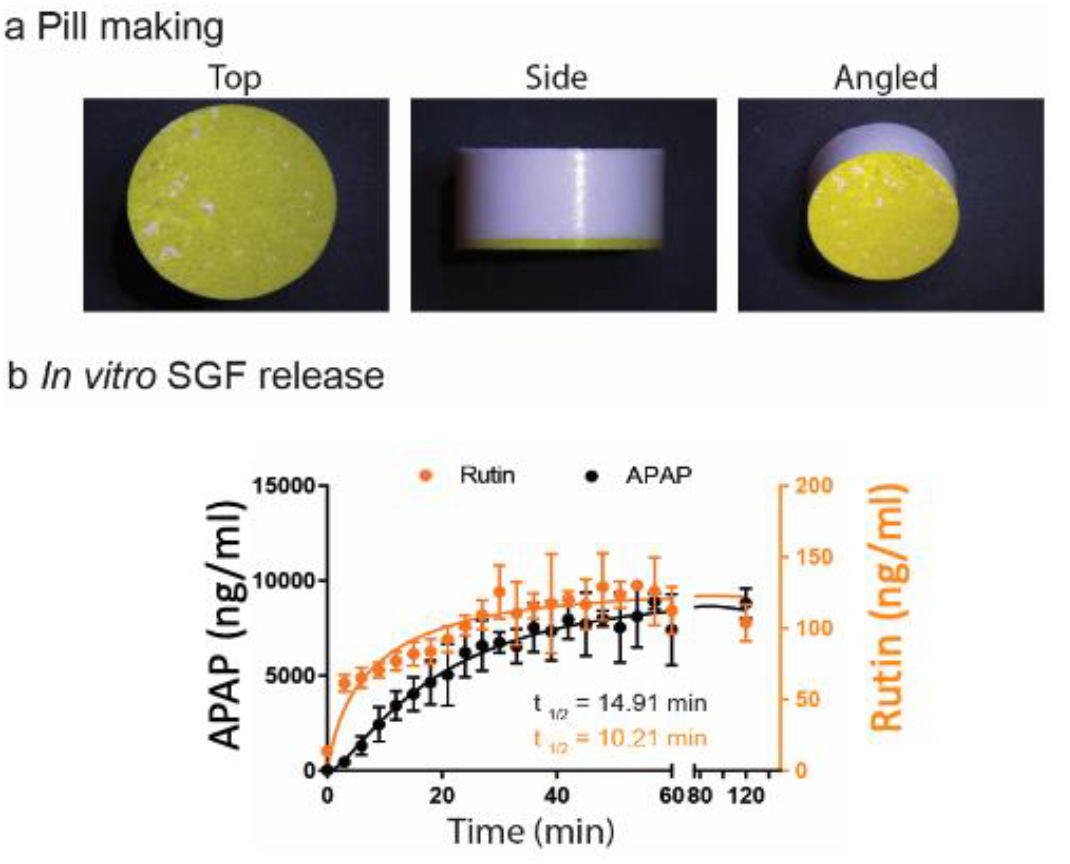
*In vitro* release profile of novel solid dosage form. A) Microscope image of novel dual formulation pill. Rutin and APAP were formulated in two separate solid formulations to enable the controlled, separate release of Rutin or APAP. The two formulations were pressed into a single 12mm pill using a two-step pill press process. B) Release profile of APAP and rutin in stimulated SGF. Rutin was in a 10-fold dilution. Concentrations of Rutin and APAP were determined by HPLC-MS. Each data point was calculated by three individual pills.

## Discussion

A detailed understanding of the metabolic profiles of small molecules and their potential interactions is critical for the development of safe and efficacious therapeutics. Adverse drug metabolism continues to be responsible for 20-40% of fulminant hepatic failure; unexpected hepatotoxicity is among the primary reasons for the withdrawal of approved drug^34-36^. Here, we report a novel experimental screening system that retains physiologically-relevant liver architecture to characterize liver drug metabolism and drug-drug interactions. Through the inclusion of machine learning methods trained on publicly-available data, we demonstrated how our platform can be used to identify functional formulations that mitigate adverse drug metabolism and the potential for drug-drug interactions. Our functional excipients were specifically selected from repositories of FDA-approved compounds; which should accelerate the development, approval, and translation of such functional formulations. Significantly, our excipients exhibited better toxicity profiles at escalated concentrations compared to previously known cytochrome inhibitors investigated here (*cf*. Figure 3a-d) and, accordingly, exhibited no adverse toxicity *in vivo*. Toxicity of APAP develops at a dose of 7.5 g/day 140 mg/Kg^37^ and we have observed a protective effect of orally applied rutin at a dose of 20 mg/Kg. Further formulation work and safety studies are required but co-administration of (12.5 % rutin m/m) is technically and economically feasible given the high medical burden of APAP intoxication. By combining technologies such as machine learning and tissue engineering, we were able to identify novel formulations that have the potential to be used for safer pharmaceutical products.

## Materials & Methods

### Tissue dissection and preparation

Porcine liver tissue was obtained from a local abattoir (Lemay & Sons, Hilltown Pork, and Blood Farm). Tissue was placed in a pre-chilled (4 ºC) UW solution (Bridge to Life Solutions LLC, Columbia, SC). The tissue was cut into slices with thicknesses of 150-250 μm using a dermatome (Zimmer Dermatome AN). The resulting slices were placed between a magnet plate for the interface system as previously described (*cf*. Figure 1)^25^ and cultured at 95% oxygen, 5% carbon dioxide, at 37°C.

### Ex vivo liver damage Live/Dead Assay

A live/dead cytotoxicity kit (Invitrogen, Molecular probes, Eugene, OR) was used to measure cell viability and proliferation during culturing. Briefly, viability was determined by the presence of intracellular esterase activity. To this end, enzymatic conversion of non-fluorescent and membrane-permeable calcein-AM (2 μM) into fluorescent calcein was tracked. Secondly, cell membrane integrity was assessed by adding ethidium homodimer-1 (EthD-1, 4 μM) which does not perfuse across membranes and therefore will only enter cells with damaged membranes, where it will bind to nucleic acid contents causing red fluorescence. After adding both signaling compounds, the cells were incubated at room temperature for 30 minutes. Fluorescence was monitored using a Fluoroskan reader (Tecan, Männedorf, Switzerland) with excitation at 485 nm and 530 nm emission wavelength.

### Tissue treatment

Tissue slices were placed between two customized 96-well magnet plates as previously described (Figure S2).^25^ Cytochrome inhibitors (Table S1) were prepared in culture medium and further diluted for dose-dependent experiments. In each well, we added 50 μL of prewarmed culture medium with different concentrations of the investigated inhibitors and subsequently incubated for 0.5h, 1h or 2h. This incubation period was followed with the addition of 50 μL APAP dissolved in culture medium (Sigma Aldrich, 10 mM) and was incubated for an additional 6h before assessing viability as described above.

### Machine learning

ChEMBL24.1 data^28^ containing small molecules modulating cytochrome P450 activity was downloaded and processed in KNIME. Briefly, compounds that had an annotated IC50/EC50 or Ki value of less than 10 uM were annotated as “active”. Compounds that had a value higher than 10uM or were annotated as “inactive”. Compounds that were annotated as “inconclusive” in the database were discarded. This resulted in 8451 inhibitors and 21369 non-inhibitors for 3a4, 6311 inhibitors and 13433 non-inhibitors for 1a2, and 22 inhibitors compounds and 936 non-inhibitors for 2e1. To augment the short list of active inhibitors for 2e1, we added manually extracted inhibitors from the literature, namely 4-methylpyrazole, 4-phenyl-5-methyl-1,2,3-Thiadiazole, 4,5-diphenyl-1,2,3-Thiadiazole, Cinnamaldehyde, Clomethiazole, Clotrimazole, Curcumin, Diallyl disulfide, Diethyldithiocarbamic acid, Dimethylformamide, Dimethyl sulfoxide, Disulfiram, Genistein, Hexane, Isoniazid, Kaempferol, Midostaurin, Myricetin, Nabilone, Nifedipine, Orphenadrine, Piperine, Pyridine, Resveratrol, Rhein, Rufinamide, Safrole, Tannic acid, Ticlopidine, Zucapsaicin. We described all molecules using Morgan fingerprints (RDKit, r=4, 2048 bits) as well as RDKit physicochemical properties.^38^ Based on this data, we build random forest machine learning models in scikit-learn^39^ with 500 trees. Performance for these models were evaluated using BEDROC values based on 10-fold cross validation. For the first prospective compound sets, small molecules in the “approved” category were extracted from the DrugBank database (version 5.0)^40^ and filtered for compounds that had a Tanimoto similarity larger than 0.5 based on Morgan fingerprints (RDKit, r=2, 2048 bits); this was to ensure novelty. The cytochrome P450 2e1 model was used to anticipate the inhibitory potential of these compounds and then the compounds were ranked according to the number of trees in the random forest model that classified the compound as an inhibitor (maximum predictive confidence). For the second prospective study, we extracted generally recognized as safe (GRAS) and inactive ingredient (IIG) compounds from the FDA website and curated the collection according to previously published protocols^15,16^. Compounds were then ranked separately by the cytochrome P450 1a2 and 3a4 models according to maximum predictive confidence. A final prediction rank for each candidate compound was determined by averaging the 1a2 rank and the 3a4 rank for a compound, thereby balancing predictive confidence for a compound to be a cytochrome 1a2 and 3a4 inhibitor. For example, Eugenol ranked as the 6^th^ most promising inhibitor for cytochrome 1a2 and the 27^th^ most promising inhibitor for cytochrome 3a4, therefore leading to an average rank of 16.5 and making it the most promising candidate to exhibit dual-inhibitory activity among all 827 investigated excipients. The second most promising candidate Cyanocobalamin had an average rank of 17.5.

### *In vivo* experiments

6-8 weeks old, female Balb/c mice (Charles River) were housed at the Koch Institute of Integrative Cancer Research at MIT on a 12-hour light/12-hour dark cycle and supplied with normal diet (Lab Supply, 5002) *ad libitum*. Mice were fasted for 12 h prior to cytochrome inhibitor and APAP administration. Rutin and eugenol were prepared in PEG400 and sterile filtered (Millipore, Millex-sp 0.22 μm sterilized filter) prior to delivery *via* oral gavage.

APAP was prepared in PBS at a concentration of 20 mg/ml and sterile filtered (0.2 μm filter) prior to intraperitoneal (IP) injection at dosages of either 350 mg/kg (toxic level) or 100 mg/kg (safe level). ALT and AST plasma levels of treated mice were determined 24h after treatment through the IDEXX Chem 21 SDMA test. Additionally, intact liver was dissected and fixed using a 4% PFA for histological processing. A separate sample was snap frozen for lipid oxidation, superoxide, and total GSH concentration assays as well as cytochrome P450 expression analysis by qPCR. Specifically, the lipid oxidation and superoxide level assays were conducted according to manufacturer’s specifications (Lipid peroxidation assay kit, Sigma Aldrich Cat#MAK085; Fluorimtric hydrogen peroxide assay kit, Sigma Aldrich Cat#MAK165). Total GSH level of liver tissue was measured using a glutathione colorimetric detection kit (ThermoFisher Scientific, Cat#EIAGSHC) according to the manufacturer’s protocol. Level of GSH was normalized by the total protein amount (BCA assay, Pierce Cat#23227) of each sample to account for variations in tissue amounts.

### Pharmacokinetic evaluation of acetaminophen, major metabolites and liver

From the *in vivo* experiment, we were able to determine the lowest concentrations of rutin, eugenol and diallysulfide (i.e., DAS) that provided hepatotoxicity protection; these concentrations were used to determine how the inhibition of CYPs would affect the pharmacokinetics of APAP and its major metabolites. Inhibitors were orally administrated to female Balb/c mice followed by intraparietal injection of toxic (high) and safe (low) dose of APAP (i.e., 350 mg/kg and 100 mg/kg animal weight respectively). For each time point (i.e., 5, 10, 15, 30, 45, 60, 120, and 240 mins), blood was sampled from five mice and serum was analyzed though a Waters ACQUITY UPLC®-I-Class System aligned with a Waters Xevo® TQ-S mass spectrometer (Waters Corporation, Milford MA). Stock solutions were prepared in methanol at a concentration of 1mg/mL. A twelve-point calibration curve was prepared in analyte-free, blank mouse serum ranging from 1.25-60000 ng/mL. 20 μl of each serum sample was spiked with 40 μl of the relevant internal standard in acetonitrile at 250 ng/mL to elicit protein precipitation. APAP d4 and 4-Acetamidophenyl-β-D-Glucuronide d3 were used as the APAP internal standards in positive and negative ionization modes respectively. Thymol was the internal standard for eugenol, and tolbutamide was the internal standard for rutin. Since eugenol is nonpolar and does not contain an ionizable group, a simple derivatization reaction with dansyl chloride was performed based on an established protocol^41^. The detailed sample preparation and analysis is described in Supporting Information.

### Pill formulation and Pill press

The CYPs inhibitor formulated APAP pill was made of two active formulation components (i.e., APAP and rutin powder) and relative excipients to control the release of active drugs. All ingredients were grinded before pill pressing in order to make fine powders. Two layers were composed for each tablet, i.e., the APAP layer and the CYPs inhibitor layer. For the APAP layer, 500 mg APAP, 72 mg carboxyl cellulose, cellulose (216 mg), and 72 mg magnesium stearate were mixed and pressed under 8000 N with a 12 mm dice compression force pill pressor (Natoli NP-RD10A). For the CYPs inhibitor layer, 58 mg of rutin, 7 mg of sodium bicarbonate and 7 mg citric acid were mixed and pressed on top of the APAP layer under 26000 N with the same dice.

### *In vitro* pill release

A standard dissolution test for our pressed tablets was performed to determine release kinetics of APAP as well as rutin on a Hanson Vision Elite according to manufacturer protocols. Briefly, three tablets were submerged separately in three 800 mL containers filled with synthetic SGF (pH 2.5). The dissolution experiment was performed while constantly stirring each of the solutions at 75 rpm. The solutions were sampled using the integrated autosampler every 3 minutes by extracting 1ml of sample from each chamber and deposited it into separate glass vials. After the experiment was complete, all samples were prepared for high-performance liquid chromatography in order to analyze molecular concentration (*cf*. Supporting Methods and Materials). APAP and rutin concentrations were averaged for the three individual experiments and plotted over time. A Gaussian curve was fitted in order to calculate the half-life of the release (t_½_).

### Statistical Analysis

All experiments were performed at least three times, and we present the data as the mean with standard deviation. An analysis of variance (ANOVA) was performed using Prism 7 (Graphpad, San Diego, Calif., USA) and a value of p < 0.05 was considered statistically significant and was noted in all figures, the significance at p < 0.05, 0.01 and 0.001 were labeled as *, ** and ***, respectively.

**Table 1.**
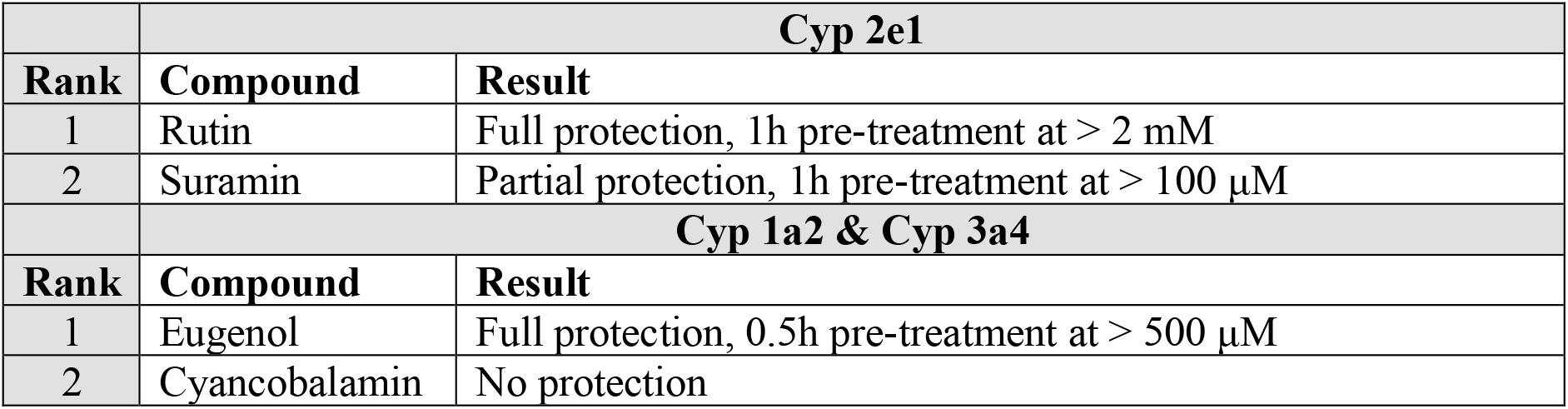
Summary of the *in vivo* protection effect of the top rank molecules predicted from in-house Cyp2e1, 1a2 and 3a4 substrate database from APAP induced liver toxicity.

| Cyp 2e1 |  |  |
| --- | --- | --- |
| Rank | Compound | Result |
| 1 | Rutin | Full protection, 1h pre-treatment at > 2 mM |
| 2 | Suramin | Partial protection, 1h pre-treatment at > 100 μM |
| Cyp 1a2 & Cyp 3a4 |  |  |
| Rank | Compound | Result |
| 1 | Eugenol | Full protection, 0.5h pre-treatment at > 500 μM |
| 2 | Cyancobalamin | No protection |

## Acknowledgements

We would like to thank Prof. R. Langer for helpful discussion. The authors would like to acknowledge the use of resources of Microscopy Core Facilities at Swanson Biotechnology Center, and David H. Koch Institute for Integrative Cancer Research at MIT. The authors would also like to acknowledge partners RPDR team for their assistance with identifying patients and Dr. Miguel Jimenez for his significant contributions to editing the manuscript.

## Funding

C. L. was supported by a postdoctoral fellowship and funding from the National Science Council of Taiwan (Grant 106-2917-I-564-082 and 110-2222-E-038-001-MY3) D.R. is a Swiss National Science Foundation Fellow (Grant P2EZP3_168827 and P300P2_177833). This work was in part supported by NIH grant EB000244 (G.T.) the Karl van Tassel (1925) Career Development Professorship and Department of Mechanical Engineering MIT and Division of Gastroenterology, Brigham and Women’s Hospital (G.T.).

## Contributions

Y. S., C. L., D. R., and G. T. conceived the study, designed experiments and analyzed data. C. S., K. H., J. C., S. T., K. I., A. L., and J. W., performed experiments. Y. S., C. L., D. R., C.W., and G. T., wrote the manuscript. W. G., and G. T. supervised the study. All authors discussed the results and assisted in the preparation of the manuscript.

## Competing Interests

D.R. acts as a consultant for the pharmaceutical and biotechnology industry and a mentor for the German Accelerator Life Sciences. C. S. is currently employed by Bayer Pharmaceuticals. Complete details of all relationships for profit and not for profit for G.T. can found at the following link: https://www.dropbox.com/sh/szi7vnr4a2ajb56/AABs5N5i0q9AfT1IqIJAE-T5a?dl=0.

## Data Availability

All data associated with this study are present in the paper and/or the Supplementary Materials.

