## Supplementary_Information for "A machine learning liver-on-a-chip system for safer drug formulation"

### Supporting Information

**Yunhua Shi[1,2]†, Chih-Hsin Lin[1, 6]†, Daniel Reker[1,2, 7]†, Christoph Steiger[1,2], Kaitlyn Hess [1], Joy E. Collins[1], Siddartha Tamang[1], Keiko Ishida[1], Aaron Lopes[1,2], Jacob Wainer[1,2], Alison M. Hayward[1,2,3,8], Chad Walesky[4], Wolfram Goessling[4,5], and Giovanni Traverso*[1,2,3]**

[1] Koch Institute for Integrative Cancer Research, Massachusetts Institute of Technology, Cambridge, MA 02139 (USA)

[2] Division of Gastroenterology, Hepatology and Endoscopy, Department of Medicine, Brigham and Women's Hospital, Harvard Medical School, Boston, MA 02115, USA

[3] Department of Mechanical Engineering, Massachusetts Institute of Technology, Cambridge, MA 02139 (USA)

[4] Division of Genetics, Brigham and Women's Hospital, Harvard Medical School, Boston, MA 02115 (USA)

[5] Division of Gastroenterology, Massachusetts General Hospital, Harvard Medical School, Boston, MA 02114 (USA)

[6] Graduate Institute of Nanomedicine and Medical Engineering, College of Biomedical Engineering, Taipei Medical University, Taipei 11031, Taiwan

[7] Department of Biomedical Engineering, Duke University, Durham, NC 27708 (USA)

[8] Division of Comparative Medicine, Massachusetts Institute of Technology, Cambridge, MA 02139 (USA)

† These authors contributed equally.

**Material and Method:**

**Histology staining and morphological assessment**

Liver tissue slices were fixed in formalin for 10 minutes at room temperature. Slices were subsequently rinsed with phosphate buffer saline (PBS) and placed in a histological cassette, which was submerged in 70% ethanol for storage. The samples were evaluated by the MIT iLab histology facility using routine hematoxylin-eosin (H&E) staining and light microscopy evaluation.

**RT-PCR and qPCR**

Total RNA is extracted from liver biopsy with Total RNA isolation kit (Zymo Research) according to manufacture protocol. After extraction, total RNA is converted to cDNA with High-capacity cDNA reverse transcription Kit (ThermoFisher). RT-PCR is performed with Platinum Taq DNA polymerase (ThermoFisher) and qPCR is performed with SYBR green real-time PCR master mixes (ThermoFisher). For RT-PCR, the product is run against 1.5% agarose gel and visualized with Bio-rad gel image suit. qPCR is run through Roche Lightcycle 480 and analyzed with double delta Ct value comparison normalized to the untreated sample group. Each individual experiment are conducted with three repeats and five different samples are included in the same group for statistical analysis.

**Adenosine Triphosphate (ATP) Assay**

The ATP assay is the most common way to measure tissue viability. It measures intracellular ATP, which is required for cellular metabolism. It is a key indicator of cellular activity and has been utilized as a measure of cell viability and cytotoxicity in research and drug discovery. Each slice was weighed and homogenized in 10% trichloroacetic acid (TCA) at room temperature. The samples were then centrifuged for 10 min at 11,000 g at 4°C. The supernatant was used for measurement of intracellular ATP. The ATP determination kit (Thermo Fischer Scientific, Molecular Probes®, A22066) provided a rapid method to measure intracellular ATP. The slice homogenate supernatant 4 ul was used for the ATP kit. If cells release ATP, then in the presence of luciferase, ATP immediately reacts with the Substrate D-luciferin to produce light. The light intensity is therefore a direct measure of intracellular ATP concentration.

**AlamarBlue Cell Viability Assay**

The AlamarBlue® Cell Viability assay (Cat. no. DAL1100, Lifetechnologies) measures quantitatively the proliferation of cells. The liver slices were placed in the 96 well magnet plate with different treatments. After removing all treatments, 65 µL AlamarBlue reagent was added into each well and incubated for 3 hours at 37°C. The fluorescence is monitored using an ELISA reader (Tecan, Männedorf, Switzerland), at 560 nm with an excitation wavelength and 590 nm emission wavelength.

**Protein Content of Liver Tissue Slices**

The pellet of the homogenized samples was used to determine the protein content of the liver tissue slices. The pellet was dissolved in 200 ul of 5M NaOH for 30 mins at 37°C. The samples were diluted with 800 ul of water. The total protein content of liver tissue slice homogenate was determined using a Bio-Rad DC protein Assay (Bio-Rad, Munich, Germany), using a bovine serum albumin calibration curve. After incubation in the dark for 15 mins, the absorbance at 650 nm was measured on an ELISA reader (Tecan, Mmännedorf, Switzerland) and compared to a standard curve.

**Human hepatocyte microsomal inhibition assay:**

Cytochrome P450 inhibition assay was performed using the Vivid CYP450 screening kits (Invitrogen, CYP3A4 blue and CYP1A2 blue) according to the manufacture's protocol. Briefly, Vivid substrates are reconstituted in DMSO (10 µM BOMCC and 3 µM EOMCC). Test compounds (rutin or eugenol) were prepared in 1×vivid CYP450 reaction buffer at a stock concentration that was 2.5-fold higher than the working concentration. Both positive substrates and test compounds were diluted in the "Master Mix" provided by the kits and incubated while being protected from light for 10 minutes (22ºC). Cytochrome activity is measure by assessing fluorescence of the proprietary substrate with 415nm excitation and 460 nm emission. To quantify percentage inhibition, fluorescence of each sample was compared to the signal from Vivid substrates as provided by the manufacturer (BOMCC and EOMCC).

**Pharmacokinetics analysis with LC-MS:** Eugenol derivitazation was performed as described below: 100 µl dansyl chloride at 1mg/mL in acetone and 20 µl of .1M KOH was added to precipitated serum samples. The reaction was performed at room temperature at 15 minutes before proceeding to the next step.All samples were briefly vortexed, sonicated for 10 minutes, and centrifuged for 10 minutes at 13,000 rpm. 50 µl of supernatant was pipetted into a 96-well plate containing 50 µl of water. Finally, 1.00 µl was injected onto the UPLC-ESI-MS system for analysis.

Analyte concentrations in serum from in vivo experiments were analyzed using Ultra-Performance Liquid Chromatography-Tandem Mass Spectrometry (UPLC-MS/MS). Analysis was performed on a Waters ACQUITY UPLC®-I-Class System aligned with a Waters Xevo® TQ-S mass spectrometer (Waters Corporation, Milford MA). Liquid chromatographic separation was performed on an Acquity UPLC® HSS T3 (50mm × 2.1mm, 1.8 µm particle size) column at 50 °C. The mobile phase consisted of aqueous 0.1% formic acid, 10mM ammonium formate solution (Mobile Phase A) and acetonitrile: 10 mM ammonium formate, 0.1% formic acid solution (95:5 v/v) (Mobile Phase B). The mobile phase had a continuous flow rate of 0.5 mL/min using a time and solvent gradient composition.

For the analysis of APAP, APAP metabolites, eugenol, and rutin the initial composition, 95% Mobile Phase A, was held for 1.00 minutes, following which the composition was changed linearly to 40% Mobile Phase A until 1.50 minutes. At 2.50 minutes the composition was 25% Mobile Phase A and 75% Mobile Phase B. At 2.80 minutes the composition was 5% Mobile Phase A. This composition was held constant until 3.50 minutes, after which the composition linearly changed back to 95% Mobile Phase A. The composition was held at 95% Mobile Phase A until completion of the run, ending at 4.50 minutes, where it remained for column equilibration. The total run time was 4.50 minutes.

The ionization mode and mass to charge transitions (m/z) used to quantitate APAP, APAP metabolites, eugenol, rutin, and their respective internal standards are in the table below. Sample introduction and ionization was by electrospray ionization (ESI) in both the positive and negative ionization mode. Waters MassLynx 4.1 software was used for data acquisition and analysis.

***In vitro* pill release HPLC analysis:** High Performance Liquid Chromatography (HPLC) was used to determine the drug concentrations from all in-vitro release assays. An Agilent 1260 Infinity II HPLC system equipped with a quaternary pump, autosampler, thermostat, control module, and diode array detector was utilized as described previously. Data processing and analysis was performed using OpenLab CDS ChemStation®. Acetaminophen, Rutin, and Eugenol were separated on an Agilent Zorbax Eclipse XDB C18 analytical column 4.6 x 150 mm with 5 µm particles, maintained at 45 °C. The optimized mobile phase consisted of A: 20 mM aqueous ammonium acetate buffer (unbuffered) and B: acetonitrile. The gradient starts at 90% A and 10% B and changes to 40% A and 60% B over a 10 minute period at a flow rate of 1.00 mL/min. The method run time was 15 minutes with a 3 minute post run. The injection volume was 5 µL, and the selected ultraviolet (UV) detection wavelength was 254 nm at a bandwidth of 4.0, no reference wavelength, and an acquisition rate of 40 Hz.

**Figures and Tables:**

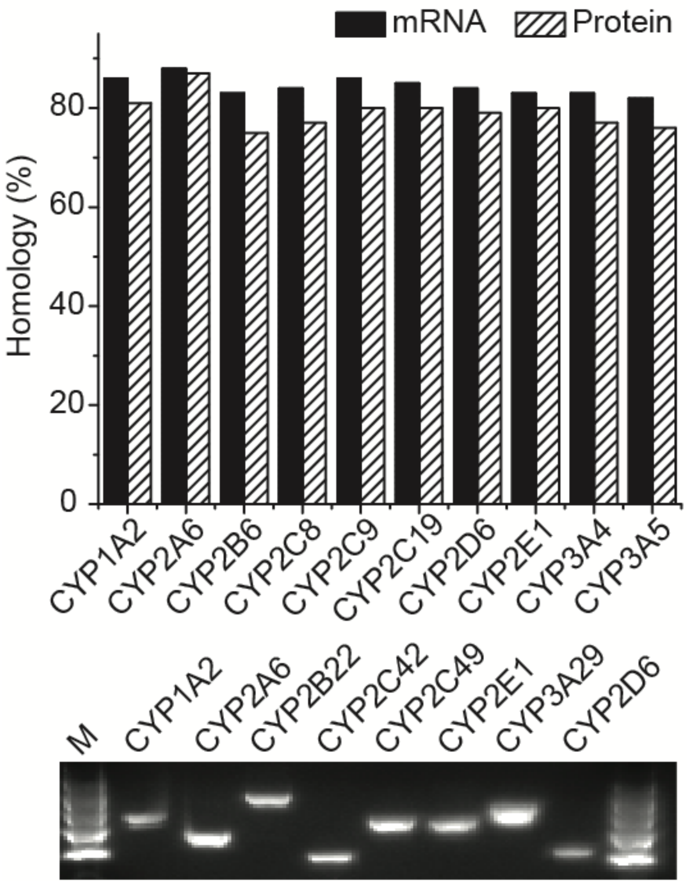


Figure S1: Homology comparison of mice, pig and human predominant liver CYPs. Upper figure, solid block represents homology comparison between human and pig and dashed block represents homology between human and mice. Bottom figure, the PCR detection of expression of major CYPs in pig liver tissue.

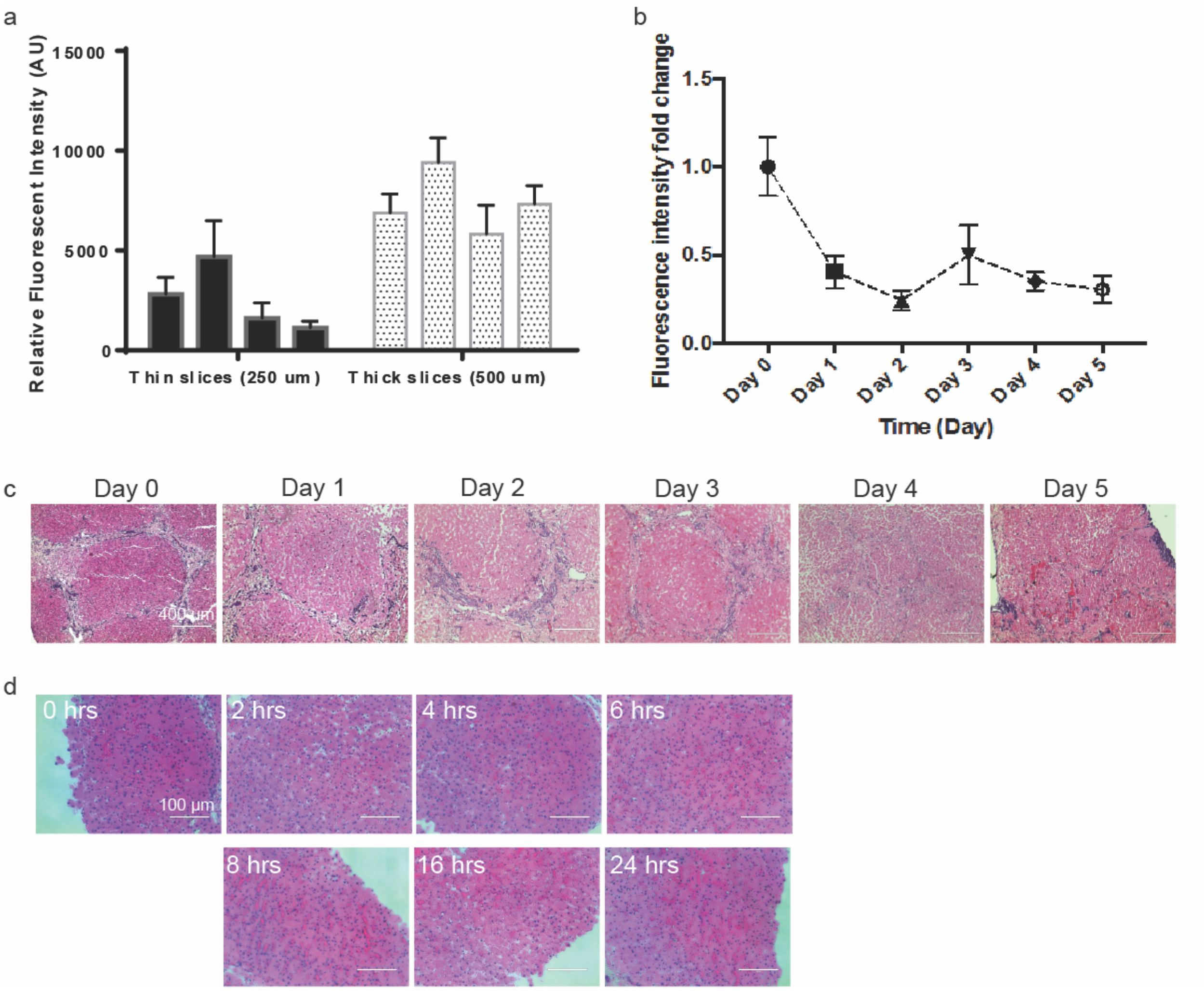


Figure S2: Platform validation additional figures. a) Live/dead assay on porcine liver slices. Liver slices with different thickness (i.e., 250 and 500 µm) were cultured for 24 hrs followed with live/dead assay. Relative fluorescence intensity indicates relative viability of cells per well. Each thickness was tested fiver times per animal, four different animal tissues (n=20) have been included to test the variability of the assay. b) *ex vivo* cultured tissue viability over 5-day time course. Viability of cell is measure with live/dead assay as described in material and method. Each time point is studied in 4 replicates for each animal and three animals are included for each point (n=12). c) H&E histology staining of liver tissue for *ex vivo* culture from Day 0 to Day 5. Bar= 400 µm. d) H&E histology staining of liver tissue for *ex vivo* culture of 0, 2, 4, 6, 8, 16 and 24 hrs. Bar= 100 µm.

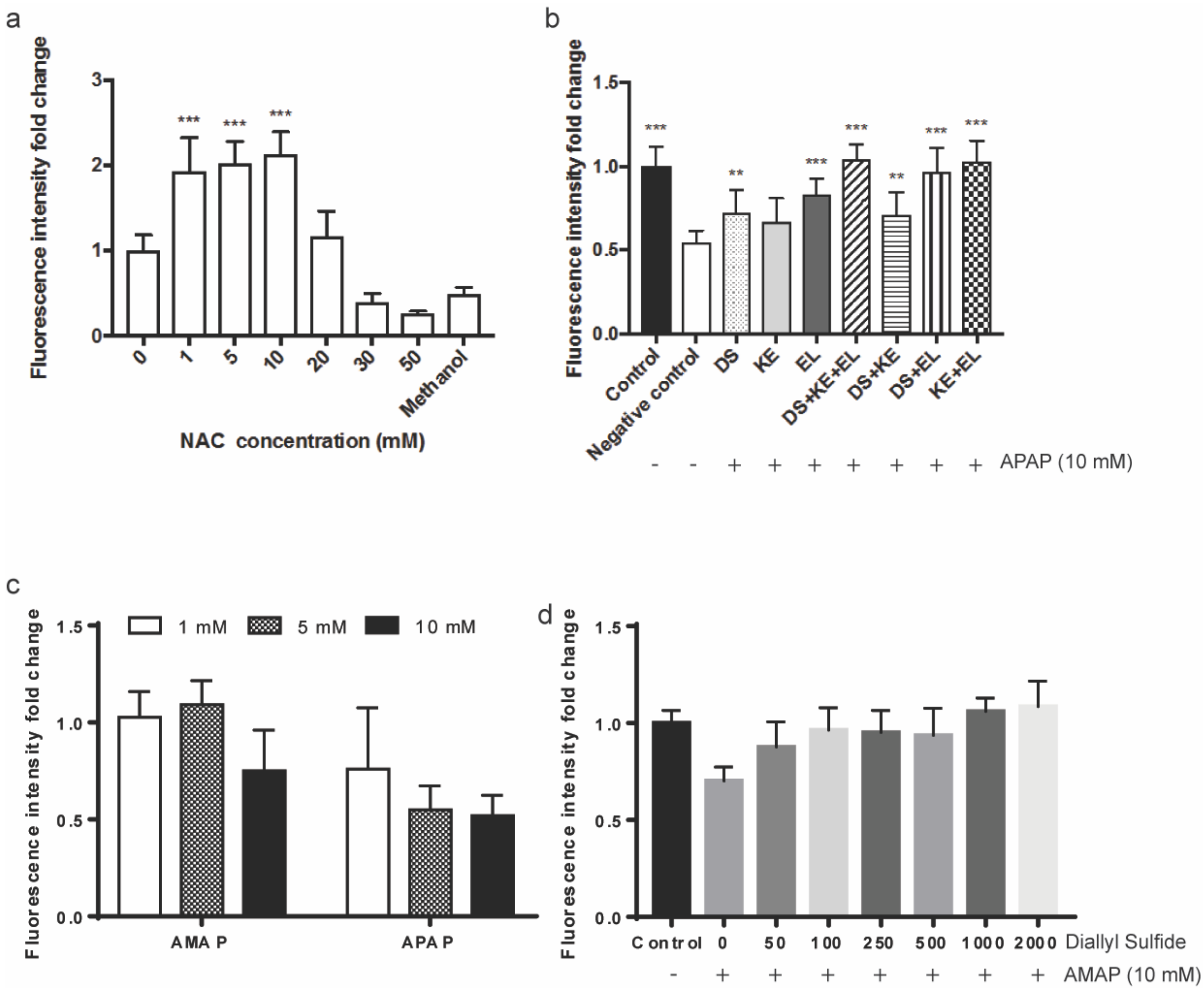


Figure S3: Platform validation additional figures

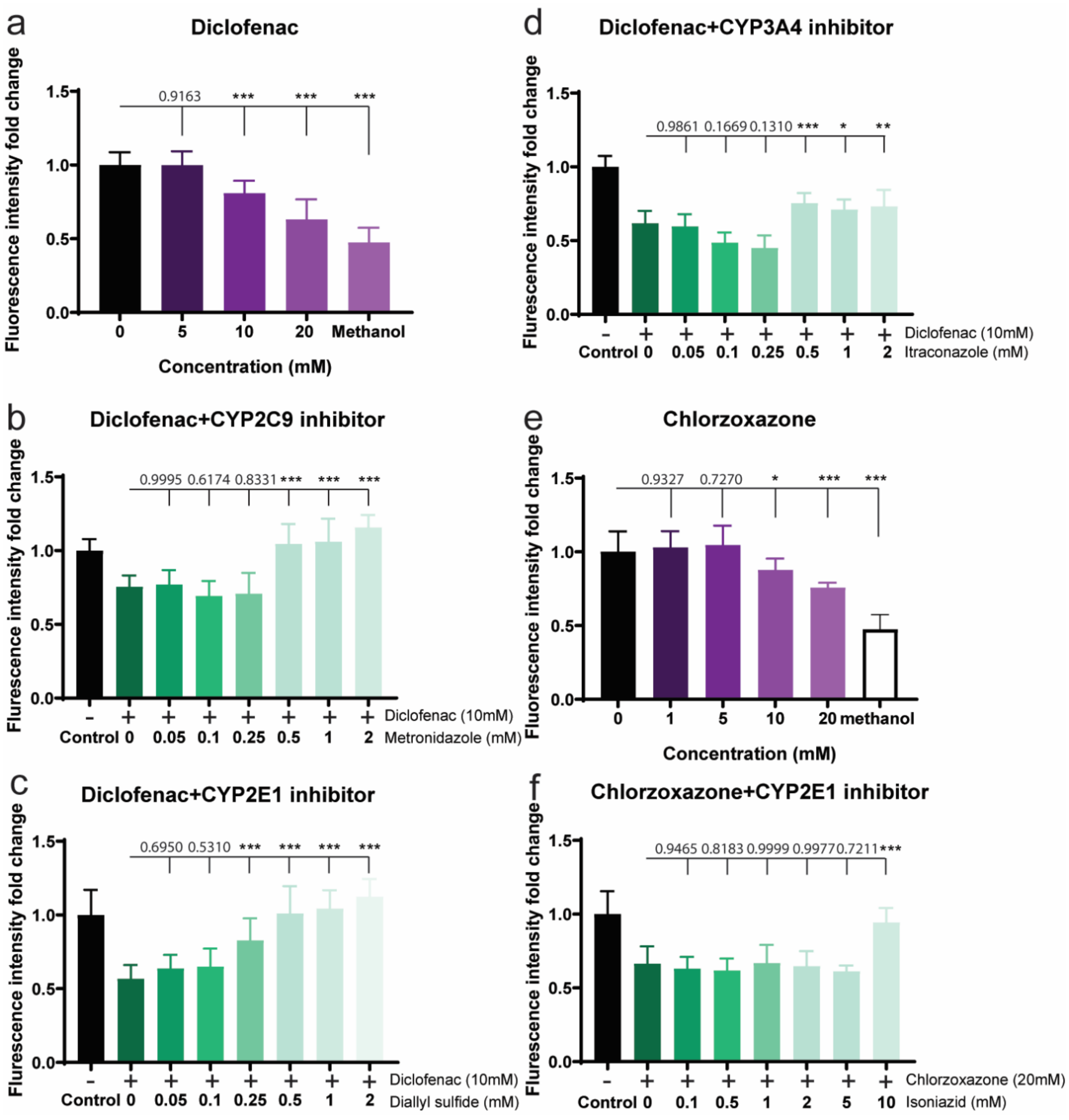


Figure S4: Platform validation with additional hepatotoxicity drugs and liver protection effect by CYPs inhibitors. a) concentration dependent hepatotoxicity of diclofenac on *ex vivo* platform. Each concentration is replicated four times on three individual animal tissues (n=12). b-d) hepatotoxicity protection effect of CYPs inhibitors (i.e., metronidazole, diallyl sulfide, and itraconazole) on liver damage induced by diclofenac (n=12). e) concentration dependent hepatotoxicity of Chlorzoxazone on *ex vivo* platform. Each concentration is replicated four times on three individual animal tissues (n=12). f) hepatotoxicity protection effect of CYP2E1 inhibitor isoniazid on liver damage induced by chlorzoxazone (n=12).

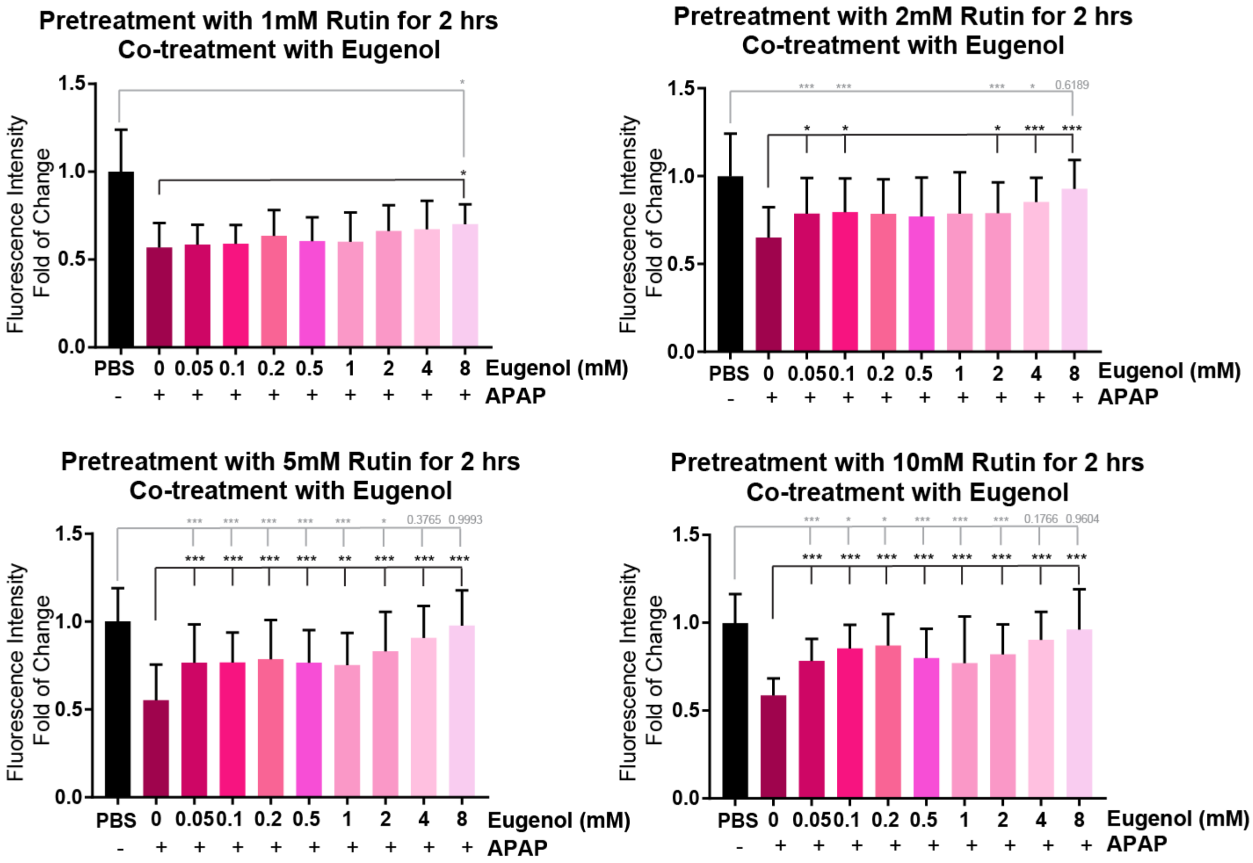


Figure S5: *Ex vivo* validation liver protection effect of the combination of eugenol and rutin in APAP overdose induced toxicity.

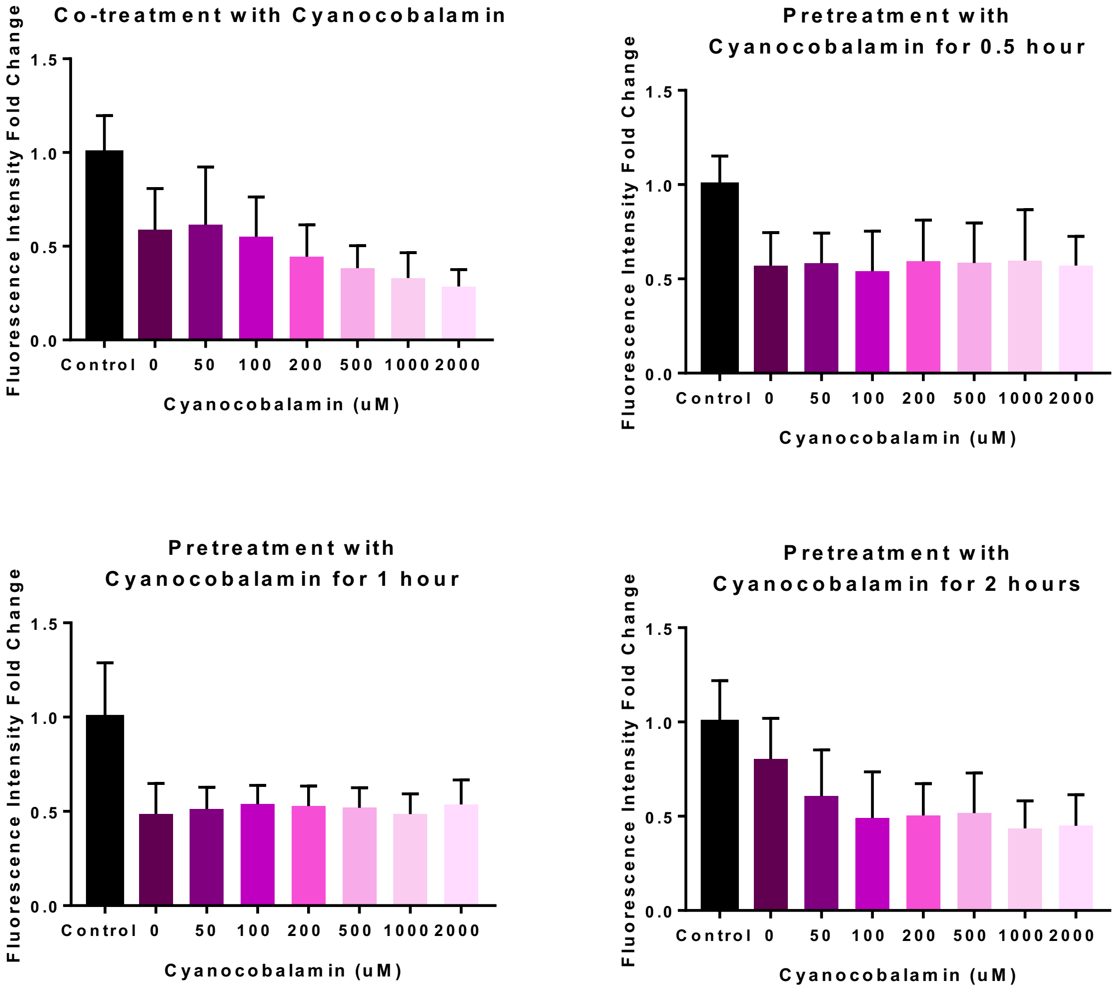


Figure S6: Ex vivo validation for cyanocobalamin.

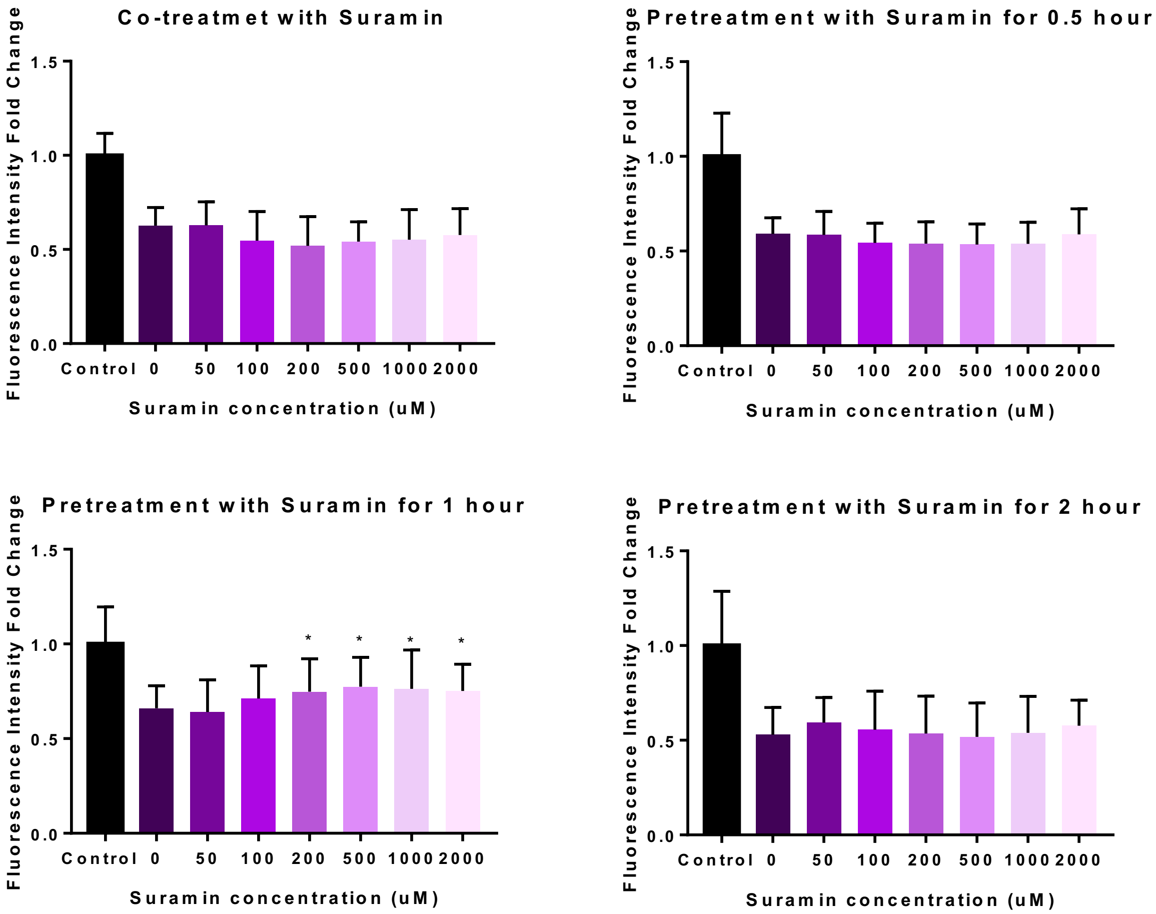


Figure S7: Ex vivo validation for suramin.

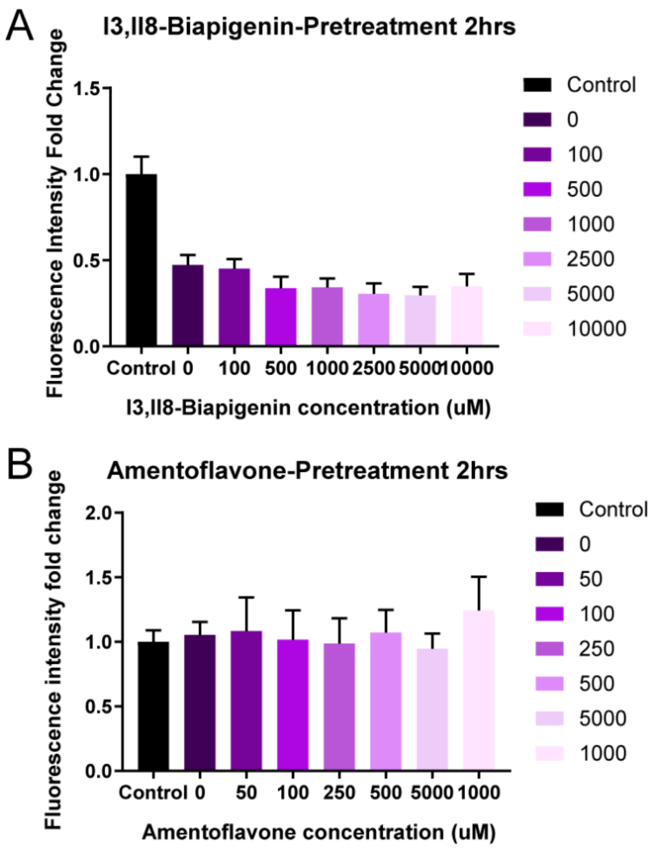


Figure S8: Ex vivo validation for 2hrs pretreatment of I3,II8-Biapigenin and amentoflavone for 10 mM APAP.

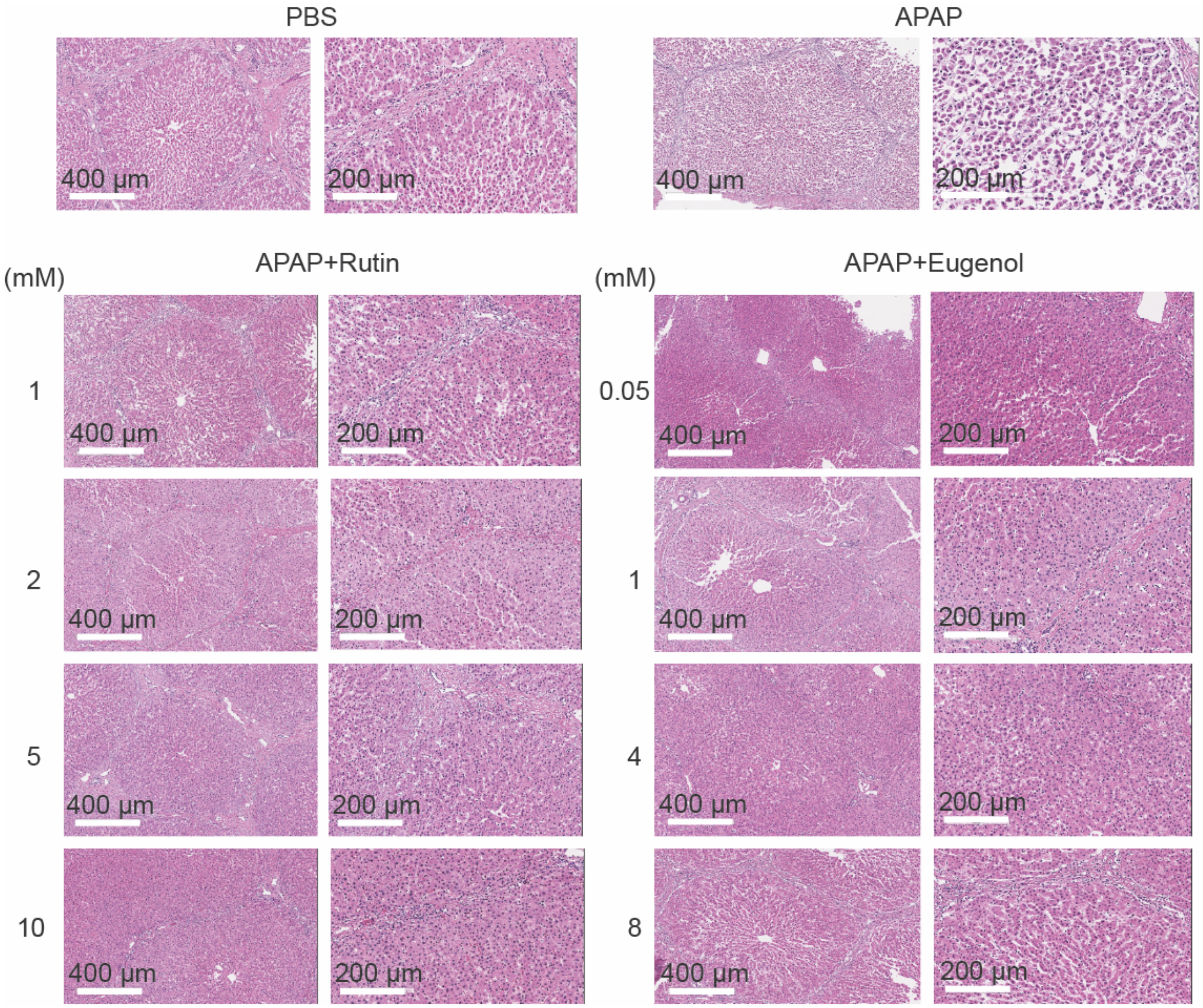


Figure S9: Histology for *ex vivo* screening tissue with rutin and eugenol administrated with APAP respectively.

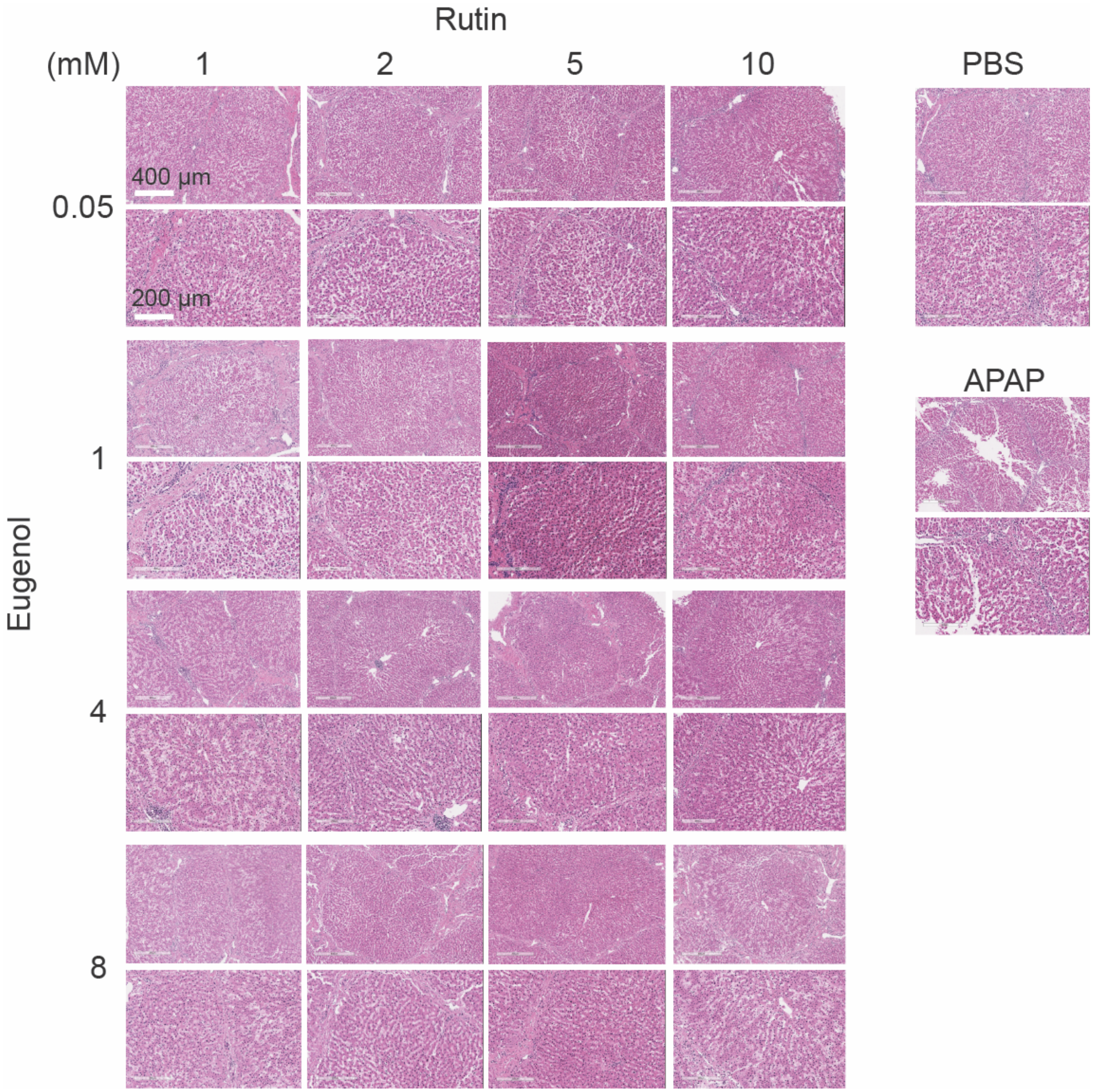


Figure S10: Histology for *ex vivo* screening tissue with rutin and eugenol together co-administrated with APAP.

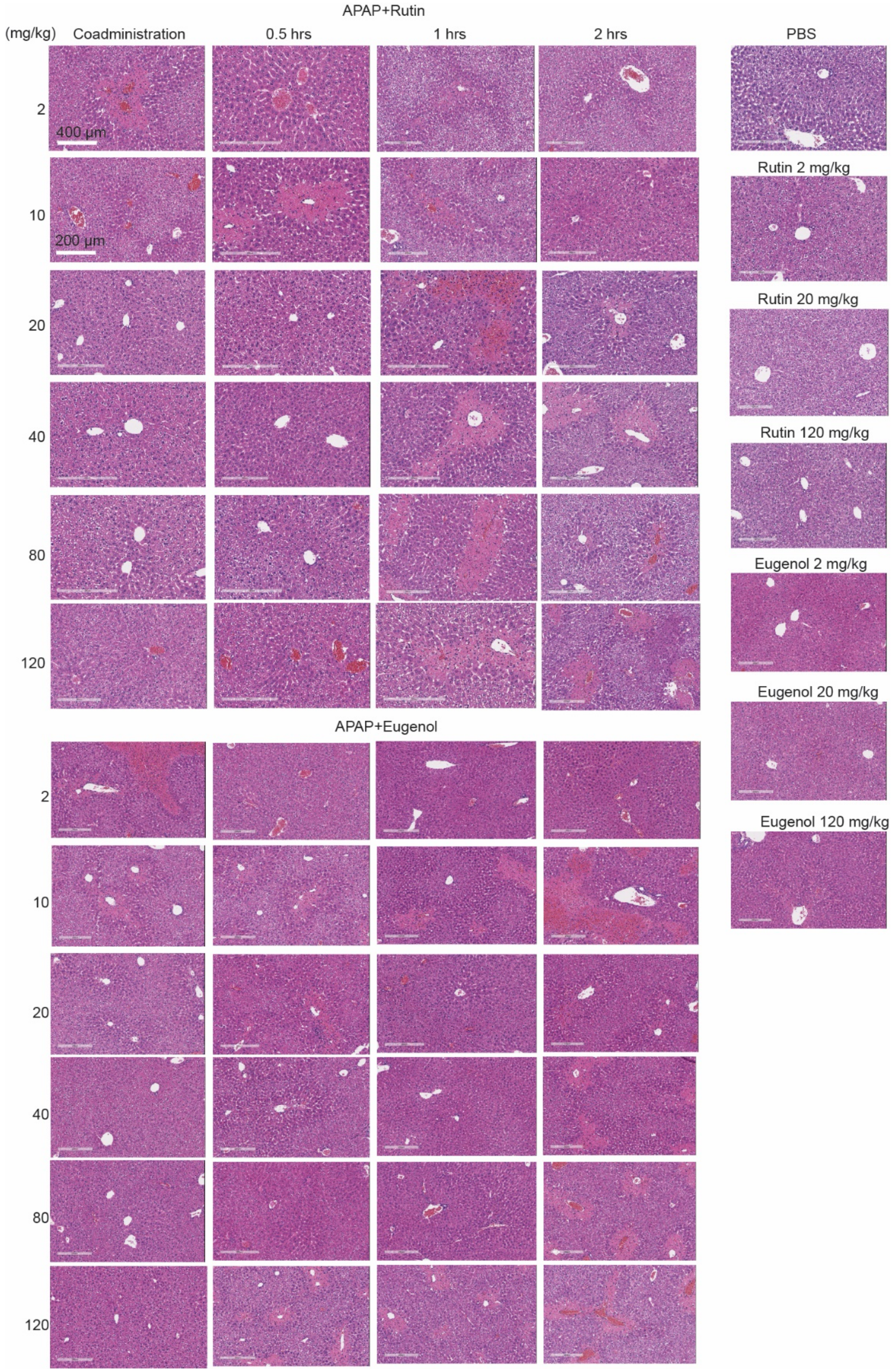


Figure S11: Histology for in vivo validation data.

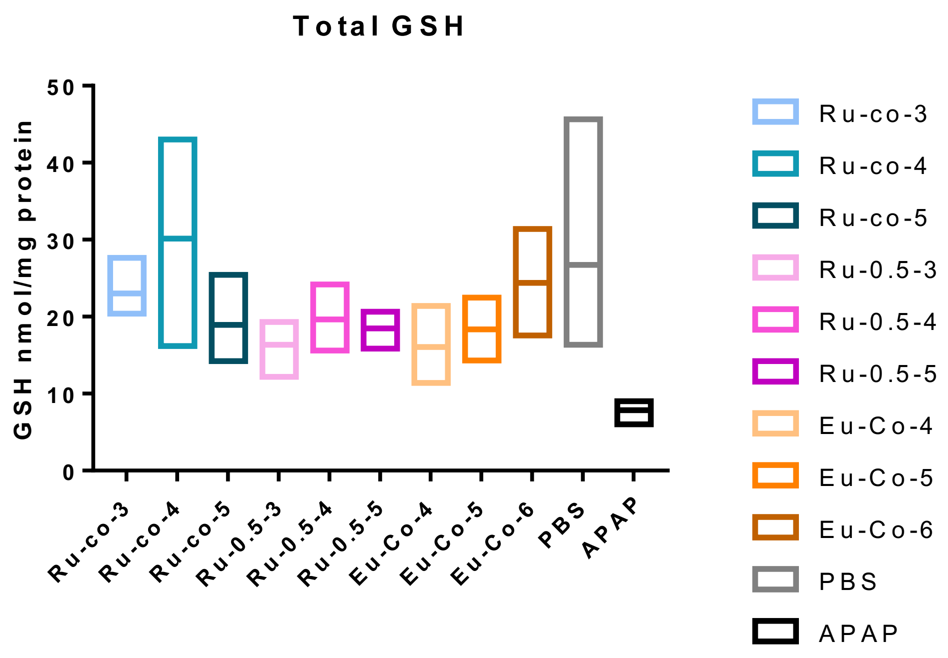


Figure S12: GSH level alteration upon rutin, eugenol and APAP administration. N=5

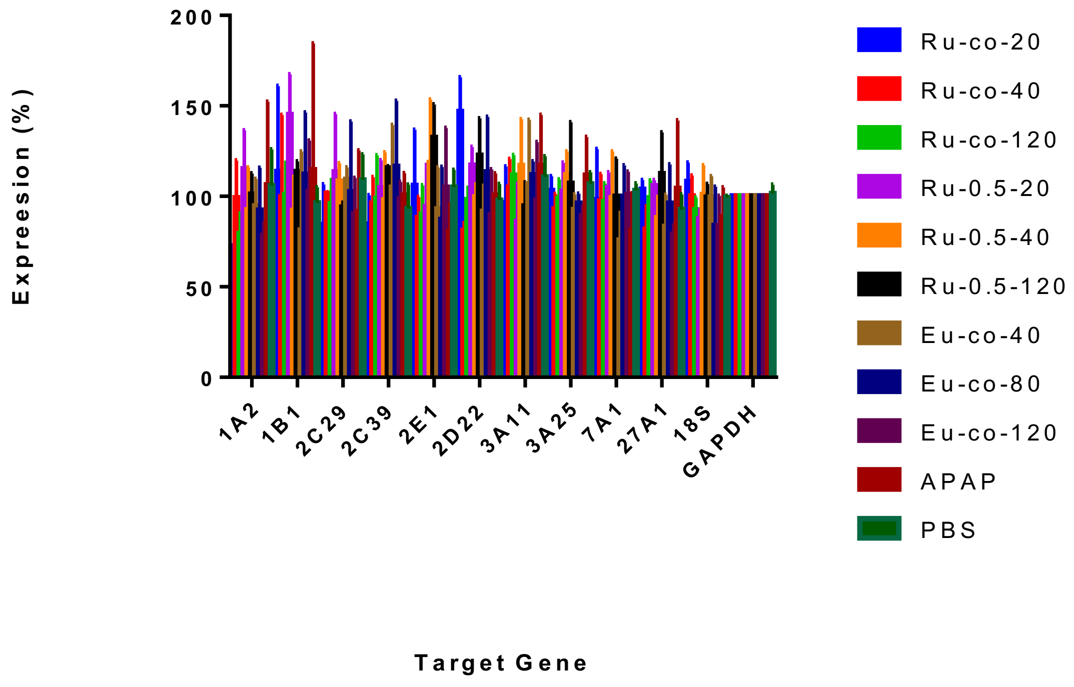


Figure S13: qPCR analysis the expression level of major CYPs in mice liver in the present of overdosed APAP, combination of rutin and APAP, and eugenol with APAP.

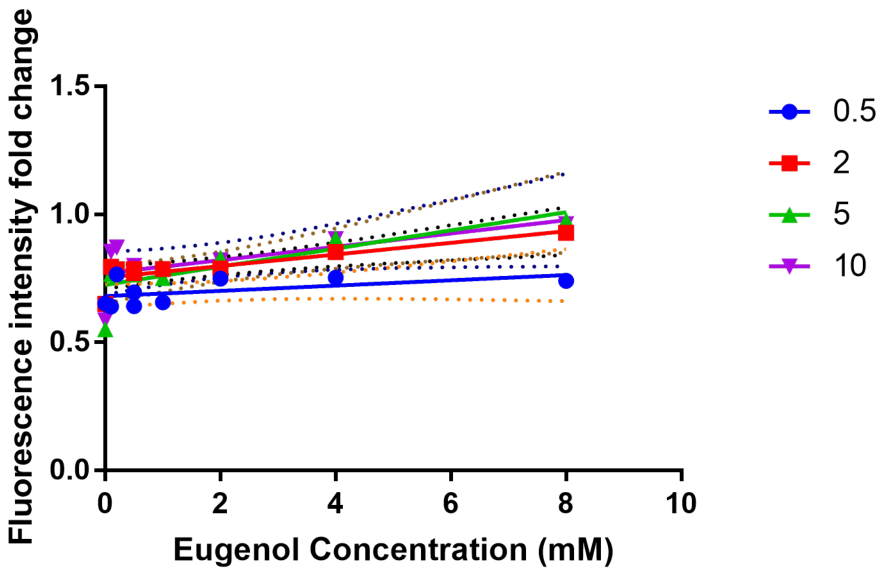


Figure S14: Lineweaver-Burk analysis of eugenol and rutin co-treatment.

| Human | | | Pig | | | Aligment (%) | |
|---|---|---|---|---|---|---|---|
| Gene | mRNA | Protein | Gene | mRNA | Protein | mRNA | Protein |
| CYP1A2 | NM_000761.4 | NP_000752.2 | CYP1A2 | NM_001159614.1 | NP_001153086.1 | 86 | 81 |
| CYP2A6 | NM_000762.5 | NP_000753.3 | CYP2A6 | NM_214417.1 | NP_999582.1 | 88 | 87 |
| CYP2B6 | NM_000767.4 | NP_000758.1 | CYP2B22 | NM_214413.1 | NP_999578.1 | 83 | 75 |
| CYP2C8 | NM_000770.3 | NP_000761.3 | CYP2C42/2C49 | NM_001167835.1/NM_214420.1 | NP_001161307.1/NP_999585.1 | 84/83 | 77/76 |
| CYP2C9 | NM_000771.3 | NP_000762.2 | CYP2C42 | NM_001167835.1 | NP_001161307.1 | 86 | 80 |
| CYP2C19 | NM_000769.3 | NP_000760.1 | CYP2C42 | NM_001167835.1 | NP_001161307.1 | 85 | 80 |
| CYP2D6 | NM_000106.5 | NP_000097.3 | CYP2D6 | KP687264.1 | AKO62672.1 | 84 | 79 |
| CYP2E1 | NM_000773.3 | NP_000764.1 | CYP2E1 | NM_214421.1 | NP_999586.1 | 83 | 80 |
| CYP3A4 | NM_001202855.2 | NP_001189784.1 | CYP3A29 | NM_214423.1 | NP_999588.1 | 83 | 77 |
| CYP3A5 | NM_000777.4 | NP_000768.1 | CYP3A29 | NM_001134824.1 | NP_999588.1 | 82 | 76 |

Table S1: NCBI notation for each liver major CYPs of pig, human and mice.

| Gene | Forward sequence | Reverse sequence |
|---|---|---|
| CYP1A2 | TCCCAGGAGAAGACCATCAA | GCCCATGCCGAAGAGAATAA |
| CYP2A6 | GATAGTGGTGCTGTGTGGATAC | CAGTGCGAGGAACTCTTTGT |
| CYP2B22 | TCCGCATGGACAAAGAGAAG | CAGAGGTCAGGATGGGATAGA |
| CYP2C42 | TAGTCTCTCCTGTCTGCTTCTC | GAGCTTCCTTCACTGCTTCATA |
| CYP2C49 | CCTCTCACTCTGGAAACAGAAC | CTCTCTCAGCCATTGGGAAAT |
| CYP2E1 | ACAAGACCTGTCTGAGGTTAATG | CATGAGAATTAGGAGCCCGTATC |
| CYP2D6 | GGAAATCGATGAGGTGATAGGG | TGACGTCAGGTTGGTGATAAG |
| CYP3A29 | CTGGCACAGATGGAGTATCTTG | GGGCAGGTAAGTGTAAGGATTT |

Table S2: RT-PCR primer used in this study.

| Gene | Forward Sequence | Reverse Sequence |
|---|---|---|
| CYP1A2 | AGTACATCTCCTTAGCCCCAG | GGGTCCGGGTGGATTCTTC |

| | | |
|---|---|---|
| CYP1B1 | CCACCAGCCTTAGTGCAGAC | GGCCAGGACGGAGAAGAGT |
| CYP2C29 | ATCTGGTCGTGTTCCTAGCG | CAGTAGGCTTTGAGCCCAAATA |
| CYP2C39 | GAGGAAGCATTCCAATGGTAGAA | TGTGAAGCGCCTAATCTCTTTC |
| CYP2e1 | CATCACCGTTGCCTTGCTTG | GGGGCAGGTTCCAACTTCT |
| CYP2d22 | TGGTTGTACTAAATGGGCTGAC | GCTAGGACTATACCTTGAGAGCG |
| CYP3a11 | GACAAACAAGCAGGGATGGAC | CCAAGCTGATTGCTAGGAGCA |
| CYP3a25 | AAGGCCATTACCATATCTGAGGA | TCTCGTCTCAAGTTTCTCACCA |
| CYP7a1 | GCTGTGGTAGTGAGCTGTTG | GTTGTCCAAAGGAGGTTCACC |
| CYP27a1 | GCACAGGAGAGTACGGAGG | CGGGCAAGTGCAGCACATA |
| 18s | AGTCCCTGCCCTTTGTACACA | GATCCGAGGGCCTCACTAAAC |
| GAPDH | AGGTCGGTGTGAACGGATTTG | TGTAGACCATGTAGTTGAGGTCA |

Table S3: qPCR primer used in this study.

| Name | Cas No. |
|---|---|
| N-Acetyl-Cysteine | 616-91-1 |
| Acetaminophen | 103-90-2 |
| N-acetyl-meta-aminophenol | 621-42-1 |
| Diallyl sulfide | 592-88-1 |
| Ketoconazole | 65277-42-1 |
| Ellipticine | 519-23-3 |
| Chlorzoxazone | 95-25-0 |
| Diclofenac | 15307-86-5 |
| Rutin | 207671-50-9 |
| Eugenol | 97-53-0 |
| Cyanocobalamin | 68-19-9 |
| Suramin | 145-63-1 |
| I3,II8-Biapigenin | 101140-06-1 |
| Amentoflavone | 1617-53-4 |

Table S4: Chemical information and preparation.
